# Protein diffusion controls how single cells respond to electric fields

**DOI:** 10.1101/2025.04.09.647627

**Authors:** Ifunanya Nwogbaga, Nathan M. Belliveau, Amit R. Singh, Daiyue Sun, Natasha Mulenga, Julie A. Theriot, Brian A. Camley

## Abstract

Cells sense and respond to electric fields, using these fields as a guidance cue in wound healing and development. This sensing is done by redistribution of charged membrane proteins on the cell’s surface (“sensors”) via electrophoresis and electroosmotic flow. If membrane proteins have to physically rearrange on the cell’s surface, how quickly can a cell respond to an applied signal? What limits the cell’s ability to respond? Are galvanotaxing cells, like chemotaxing cells, limited by stochasticity from the finite number of molecules? Here, we develop a model for the response dynamics of galvanotaxing cells and show that, for weak enough field strengths, two relevant timescales emerge: the time for the cell’s sensors to rearrange, which depends on their diffusion across the cell, and the time for the cell’s orientation to respond to an applied field, which may be very different. We fit this model to experimental measurements on the recently-identified sensor galvanin (TMEM154) in neutrophil-like HL-60 cells, finding that given the dynamics of a cell responding to an applied field, we can predict the dynamics of the cell after the field is turned off. This fit constrains the noise of the galvanotaxis process, strongly arguing that HL-60 is not limited by the stochasticity of finite sensor number. Our model also allows us to explain the effect of media viscosity on cell dynamics, and predict how cells respond to pulsed DC fields.

## I. INTRODUCTION

When epithelial tissues are injured, this disrupts the normal transepithelial potential, creating an electric field pointing toward the wound [1]. Many types of cells will polarize and respond to an applied field, migrating in the direction of the field – “galvanotaxing” or “electrotaxing” – though a few cell types also migrate opposite to the field direction [2]. While galvanotaxis is a key cue for wound healing which can override other signals [2], relatively little is understood about galvanotaxis, especially in comparison to better-characterized cell responses like chemotaxis [3]. It is generally believed that galvanotaxis requires the redistribution of charged molecules on the surface of the cell (“sensors”) via electrophoresis and electroosmotic flow [4–8], though others have argued that cells respond too quickly to an applied field for sensor motion to be essential, supporting a role for ion channels [9]. A recent CRISPR screen has identified a sensor candidate TMEM154 (“galvanin”) that is essential for galvanotaxis in HL-60 neutrophil-like cells; galvanin is redistributed to the anodal side of the cell in an applied electric field, and it is highly charged - and its charge is necessary for its function as a sensor [10]. Though there are phenomenological models of galvanotaxis [11–13] many core questions about galvanotaxis remain unanswered, including what factors limit the cell’s ability to sense the field orientation and the time required for cells to orient. In particular, for chemotaxis and chemosensing of eukaryotic cells and bacteria, there is a huge literature testing the idea that cells may be limited fundamentally by intrinsic stochasticity arising from the finite number of receptors or finite chemoattractant number [14–25], which is part of the larger field of understanding physical limits of information processing in biology [26–31]. Our earlier work developed a theory for how cells can be limited by finite sensor number [32, 33], but direct tests of this theory were impossible until the recent identification of galvanin as a sensor by Belliveau et al. [10].

In this work, we extend our theory to analytically describe the behavior of a cell exposed to a time-varying field, finding that the distribution of sensor will – for weak enough applied fields – have a linear response with a timescale limited by sensor diffusion. This weak-field limit is broadly applicable, only breaking down when the ratio of the maximum and minimum sensor concentration around the cell is ≳ 3 or so. We predict how the cell’s directionality of movement depends on the anisotropy of the sensor’s distribution around the cell’s surface. These results allow us to link cell response dynamics to the dynamics of sensor protein diffusion on the surface of the cell. We use experimental measurements of galvanin redistribution and directionality on HL-60 cells in a field to fit our model. We find good agreement with the model – but determine that finite sensor number is likely *not* the limiting factor of galvanotaxis. We also show how our model can be used to understand asymmetry between how a cell responds to a field being turned on and how a cell forgets a field after it turns off. Our results also have clear implications for technological applications of galvanotaxis, which would likely require the use of pulsed fields. Within our model, we can prove that cells in a field pulsed at x% can be no more directed on average than a field at x% strength. We fit our model to experiments measuring keratinocyte motility in pulsed fields, seeing good agreement.

## II. A LINEAR RESPONSE MODEL FOR CELL SENSOR ANISOTROPY

We begin with a simple model including sensor transport by an electric field and sensor diffusion, similar to the basic assumptions of [5, 8, 10, 32]. Sensors diffuse on the membrane with a diffusion coefficient *D* as well as having a net velocity *v*_∥_ proportional to the applied electric field parallel to the cell’s membrane, *v*_∥_ = *µE*_∥_ (*t*) (Fig. 1). The parameter *µ* can be either positive or negative – we use this parameter to encapsulate unknown details about the charge of the sensor, the details of the balance between electrophoresis and electroosmotic flow, etc.; it can be estimated from molecular details [5, 8], but here we fit it from experiment. If we assume the cell has a circular shape, the sensor position is just the angle of the sensor *θ*, and the probability distribution of sensor location *p*(*θ*) in a field *E*(*t*) pointing in the 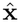 direction is then

**FIG. 1.**
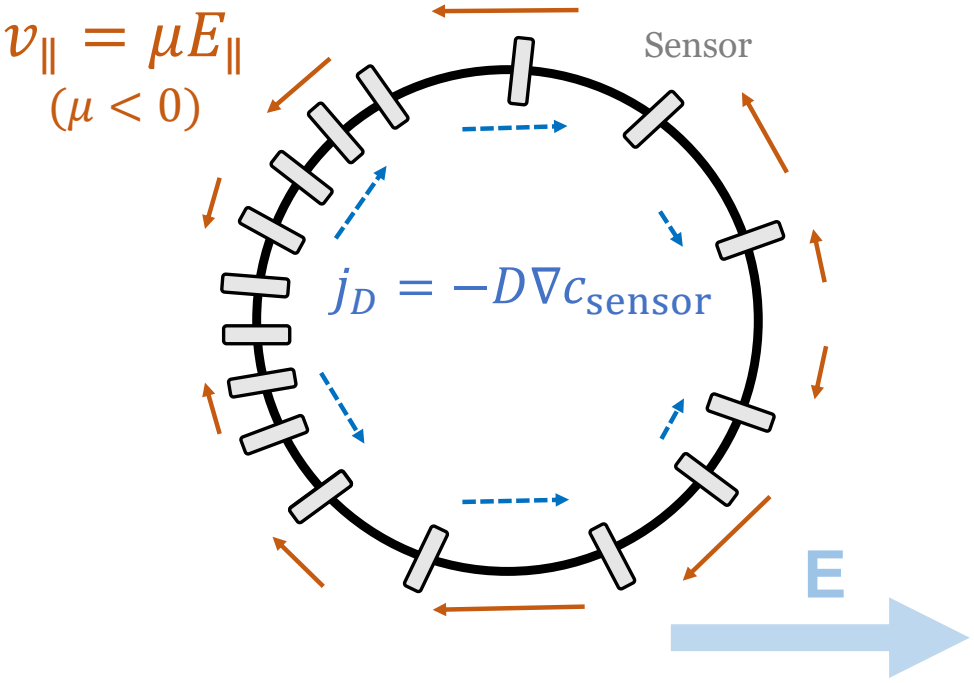
Illustration of sensors on the surface of an circular cell; sensors are labeled by their polar angle location *θ*_*i*_ and traveling with velocity *v*_∥_ = *µE*_∥_. Sensors both diffuse and undergo electromigration, leading to polarization of the sensors. We have illustrated the sensors with *µ <* 0 here, so sensors are polarized opposite to the electric field (i.e. migrate toward the anodal side of the cell)

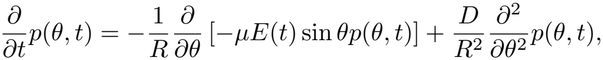

or, rescaling,

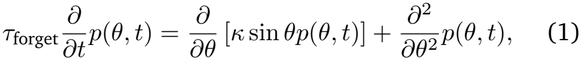

where *κ*(*t*) = *µE*(*t*)*R/D* is a rescaled electric field – essentially a Péclet number – and *τ*_forget_ = *R*^2^*/D* is the time for a sensor to diffuse the radius of the cell.

In a constant field *E*(*t*) = *E*_0_, Eq. 1 shows the sensor distribution at steady state follows a von Mises distribution

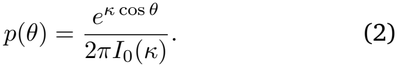

where *I*_0_ is a modified Bessel function of the first kind. We hypothesize that cells have only weak anisotropy in sensor distributions, i.e. *κ <* 1. Even if the front-back ratio of sensor concentration *p*(0)*/p*(*π*) = *e*^2*κ*^ is a factor of 2, this still corresponds to *κ* of ~0.35. If we expand to linear order in *κ*, the steady probability simplifies to

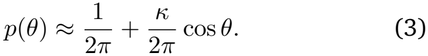

In this linear response limit, it is also possible to solve the time-response of Eq. 1 for an electric field of varying magnitude *E*(*t*) or orientation *ψ*(*t*). We start with the case where only the magnitude *E*(*t*) varies. (Oscillating orientation of the field is treated in Appendix E.) We start by writing *p*(*θ, t*) in the form 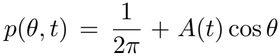, where the *anisotropy A*(*t*) is of the order *κ*_0_, with *κ*_0_ the maximum value of *κ*(*t*). If we insert this guess into Eq. (1) and neglect the term that is of order 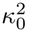, we find

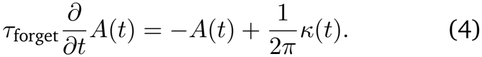

Solving using an integrating factor,

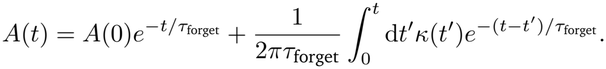

We will often work with examples where *t* ≫ *τ*_forget_ – i.e. we’ve studied the cell for long enough that the initial condition does not matter. In this case, the term proportional to *A*(0) can be neglected.

This linear response theory can be specialized to a few useful cases. For instance, if we have a cell that equilibrates initially in the absence of an electric field, and the field is then turned on to a strength *E*_0_ at a time *t*_on_, we find that the anisotropy *A*(*t*) is simple,

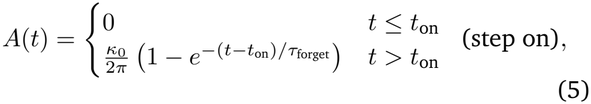

and if we have a cell that initially is equilibrated to a field of strength *E*_0_, which is turned off at a time *t*_off_, we get

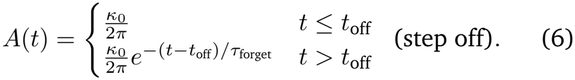

Comparing Eq. 5 and Eq. 6, we see that *the timescale for the anisotropy to be established is the same as the timescale for the anisotropy to be forgotten* – both are *τ*_forget_. This result, while mathematically unsurprising given the form of Eq. 4, is initially physically unintuitive.

We first expected that a stronger electric field would lead to a faster establishment of sensor anisotropy, as predicted by [4]. However, at weak fields, we see response time will not depend on the applied field – it is a general feature of linear response theory that the response function depends only on the unperturbed system [34]. A useful analogy is to think about an overdamped spring with a force applied to it. If we write a force balance for a spring in 1D, we get 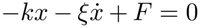. In the absence of force, the spring will decay from its original position to *x* = 0 with a timescale *τ* ≡*ξ/k*. However, with an applied force, the spring will instead relax to a new position *x*_0_ = *F/k* – but it will relax to that position with the same timescale *τ*. This is exactly the same mathematics leading to Eq. 5. Physically, we should think of the applied electric field as slightly shifting the steady-state of the cell’s sensor distribution from *A≠*0 (unpolarized) to *A* = 0 – so the time scale is the time scale of relaxation of the sensors, i.e. the time to diffuse around the cell *τ*_forget_ = *R*^2^*/D*. Ref. [35] also noted without proof that the response time and forgetting time should be equal for protein electrophoresis at weak fields, though without developing our linear response theory.

## III. CELL DIRECTIONALITIES ARE A NONLINEAR FUNCTION OF SENSOR ANISOTROPY

Given the sensor anisotropy *A*(*t*), what direction does the cell actually go? We assumed in our earlier work [32] that, given the configuration of sensors, the cell chooses an orientation that maximizes the likelihood of observing that configuration. This takes on the simple rule that the cell travels in the direction of the “vector sum” over the positions of the *N* sensors ***ρ*** = ∑ _*i*_(cos *θ*_*i*_, sin *θ*_*i*_). (Or in the opposite of this direction, if *µ <* 0 and sensors move to the back of the cell.) If we assume that the sensors are moving independently from one another, then the components of ***ρ***, 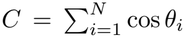 and 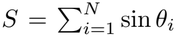, are sums of independent random variables, and we expect them to be normally distributed, with a variance *N* times the variance of each individual term, i.e. ⟨*C*^2^⟩ − ⟨*C*⟩^2^ = *N* [⟨cos^2^ *θ*⟩ − ⟨cos *θ*⟩^2^] and ⟨*S*^2^⟩ − ⟨*S*⟩^2^ = *N* [⟨sin^2^ *θ*⟩ − ⟨sin *θ*⟩^2^]. If we assume the linear response form, 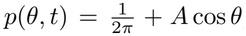, the means and variances of *C* and *S* are

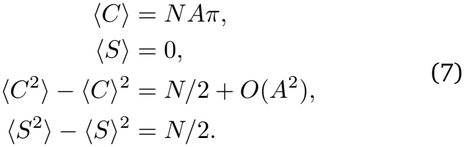

(We work this out in more detail in Appendix A.)

In the linear response limit, *C* and *S* are normally distributed with standard deviation 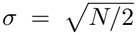, and the mean of *S* is zero. In this case the directionality (average of the cosine of the cell’s orientation) *D* can be calculated

As

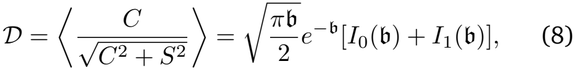

where *b* = ⟨*C*⟩ ^2^*/*4*σ*^2^ = *NA*^2^*π*^2^*/*2 is a signal-to-noise ratio and *I*_*n*_(*x*) are the modified Bessel functions of the first kind. Ref. [36] introduced this formula to study chemo-taxis; a brief derivation is given in Appendix B. This function also should in principle be multiplied by a function *ω* = sign(*A*)sign(*µ*) (Appendix B); *ω* = 1 throughout this paper, so we omit it elsewhere.

We now have a complete theory for the directionality for a cell in an arbitrary time-varying electric field:

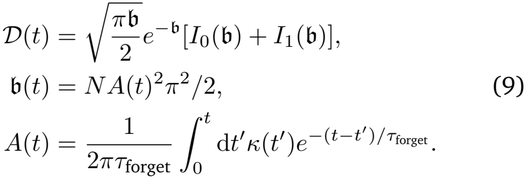

The result of Eq. 9 implies that the directionality only depends on the anisotropy through the combination of *N* and *A* in the form of *NA*^2^, which is in agreement with our bound in [32]. Therefore, scaling up the electric field strength by a factor *α* or scaling up the number of sensors by a factor *α*^2^ will have identical effects on the directionality. For this reason, we define a scale of the electric field of 1*/γ* [32] so that 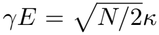. In this way, two cells with the same *γE* in a constant field will have the same directionality. While both *κ* and *γE* reflect the effective strength of the electric field, *κ* indicates how much the sensors are polarized (Eq. 2), and *γE* indicates to what extent that sensor polarization creates a directional migration.

Generally, we plot our examples assuming *κ >* 0, since this is the convention we used in [32, 33]. However, in the example of galvanin for HL-60, we find that *κ* is actually negative; plots like those shown in Fig. 2, where we rescale by *κ*, will be the same for either sign of *κ*.

**FIG. 2.**
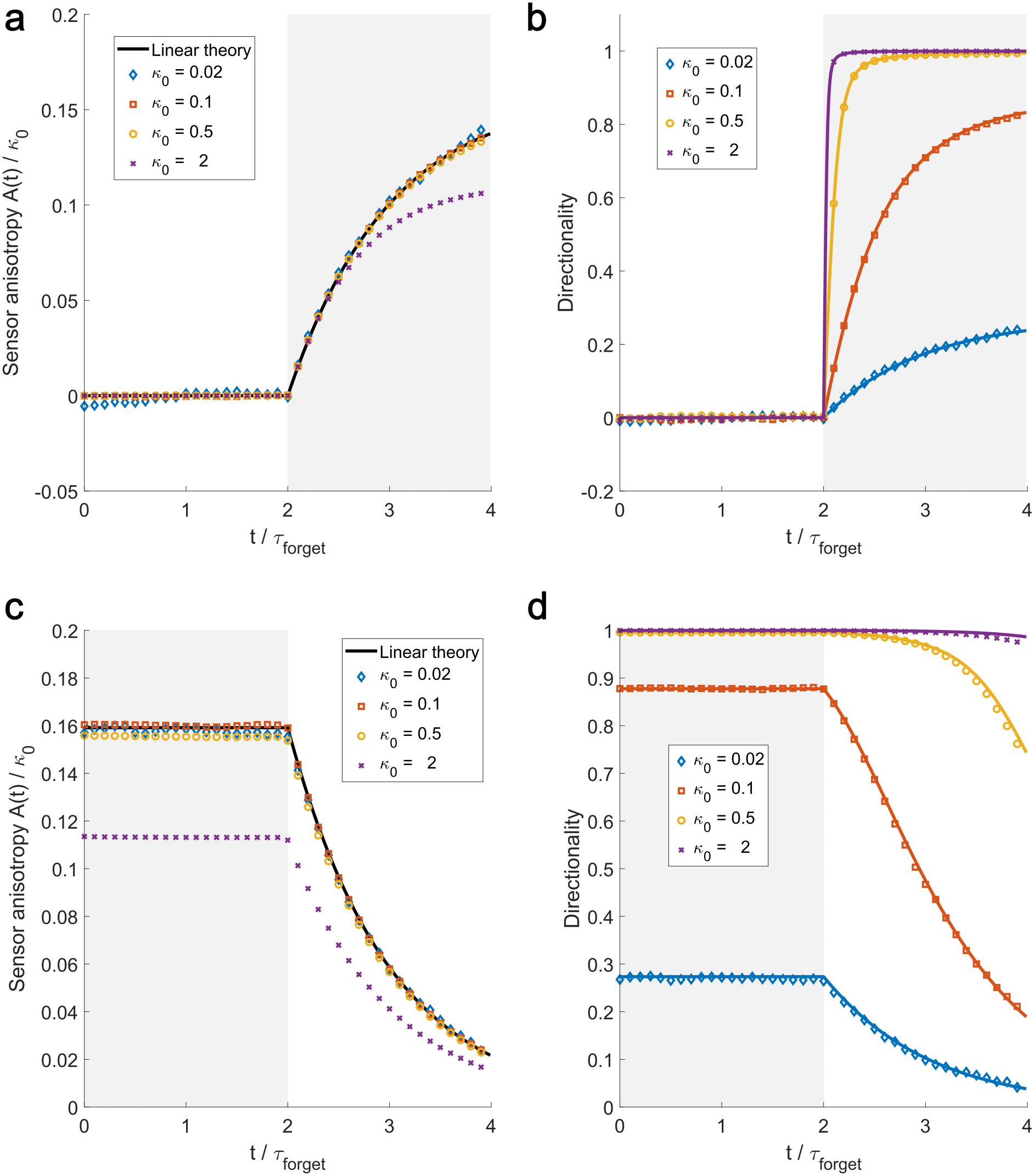
At weak field (low *κ*_0_), the response to a field being turned on or a field being turned off is completely captured by the linear theory with a timescale of *τ*_forget_, and the plots of *A*(*t*)*/κ*_0_ all collapse. However, as the field strength is increased, the cell’s directionality *D*(*t*) turns on rapidly (b) and takes a longer time to be forgotten (d). Both of these features occur because at large |*κ*|, the cell can have enough information to make a very good estimate of the field’s direction even if the anisotropy *A*(*t*) is not at its maximum value. Solid lines are theory, symbols are stochastic simulations (Brownian dynamics of diffusing and electromigrating sensors as in [32]; see Section IX.). The theory line in (a) is Eq. 5, line in (c) is Eq. 6. The solid lines in (b) and (d) are the directionality function of Eq. 9 with the appropriate *A*(*t*) curves from Eq. 5 and 6, respectively. *N* = 1000 sensors in all of these simulations. These are averaged over 5000 simulations. *A*(*t*) is extracted by constructing the histogram *p*(*θ, t*) and then computing 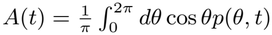.

### A. Additonal sources of noise can change relationship between anisotropy and directionality

In our discussion above, we assumed that the primary source of noise is the finite number of sensors – i.e., assuming that cells are sensing nearly at their optimum accuracy [14]. However, our results can be modified to address the possibility that either there is another source of noise in estimating the direction, e.g. stochasticity in signal transduction downstream of the sensor redistribution

[37] or if cells travel in a direction that is systematically different from the estimated direction.

To model additional stochasticity in signal transduction, instead of assuming that the cell travels in the direction 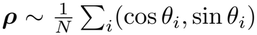, we hypothesize that the cell travels in a direction

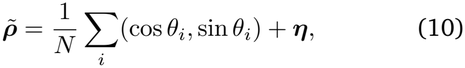

where ***η*** is a vector of two independent Gaussian noises, each with standard deviation *σ*_added_. Given this assumption for the direction of the cell, we show in Appendix A that our previous results hold in a modified form of Eq. 9 but using

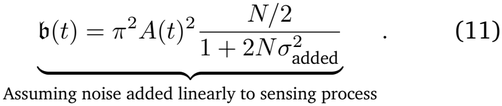

This is essentially Eq. 9 but with an effective 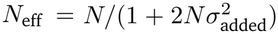. We can see that in the limit of large *N*, where the shot noise from finite sensor number is negligible, 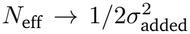. In our approach, if we fit experimental data to Eq. 9 and find a surprisingly low value for *N* relative to that expected for typical concentrations of surface molecules, we should interpret this value as *N*_eff_.

We note that, even with this additional noise, as the anisotropy *A* is increased, the effective signal-to-noise ratio *b* still increases, and directionality *D* will saturate to one. However, even if the cell “knows” the correct direction, it may not travel in that direction. For instance, keratocytes will oscillate around the electric field direction [4, 12, 38], which would lead to deviations from perfect directionality even in a deterministic model. A simple implementation of this sort of additional noise would be to assume that if the cell’s estimation of the field points it in the direction ***ρ*** with some angle Θ, the cell, due to some stochasticity in motility or oscillation, travels in a direction Θ+Δ, where Δ is uncorrelated with Θ and has mean zero. In this case then the directionality is *D* = ⟨cos(Θ + Δ)⟩ = ⟨cosΘcosΔ⟩ − ⟨sinΘsinΔ⟩ = ⟨cosΘ⟩⟨cosΔ⟩[32]. In other words, noise that rotates the cell’s orientation reduces the maximum directionality to *D*_max_ = ⟨cosΔ⟩. Thus, instead of Eq. 9, we will occasionally assume

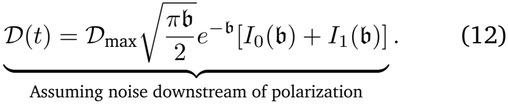

In this case, the directionality will saturate to *D*_max_ at large field strength. To distinguish this model from the one assumed in Eq. 11, we emphasize it is some noise that happens downstream of the cell’s choice of a polarization axis – i.e. the sensing error disrupts the initial choice of the cell orientation Θ but the downstream-of-polarization noise disrupts the connection between the cell’s polarization and its actual motion. Both types of noise can exist in a given cell type.

## IV. RESPONSE AND FORGETTING CURVES FOR DIRECTIONALITY APPEAR TO DEPEND ON FIELD STRENGTH

Galvanotaxis is often quantified in terms of measuring how quickly a cell responds to an applied field – the “response time” – and how quickly the cell’s directionality returns to zero after an applied field is turned off – the “forgetting time” [4, 11, 12, 39]. How should these response functions depend on field strength? Allen et al. [4] argue that response time should depend on field strength, and measure differences in the response of the directionality when field strength is varied. They also find that the response time is systematically shorter than the forgetting time. We show our theory’s predictions for response to a field being turned off or a field being turned on (and comparison to stochastic simulation) in Fig. 2. We do see that larger field strength (larger *κ*_0_) leads to a faster response *of the directionality* (Fig. 2b). Similarly, cells exposed to larger fields take a longer time to lose their directionality (Fig. 2d). However, fundamentally the *anisotropy* of the cell, the distribution of sensors, has the same dependence on time, controlled only by *τ*_forget_. *κ*_0_ sets the size of the sensor anisotropy but does not change its dynamics. We plot in Fig. 2ac the anisotropy *A*(*t*) scaled by *κ*_0_ and see that for small enough *κ*_0_, the anisotropy *A*(*t*)*/κ*_0_ collapses onto a single universal curve given by Eq. 5 for Fig. 2a and Eq. 6 for Fig. 2c.

Our analytic solution Eq. 9 says that the anisotropy of the sensors – the extent to which the sensors are polarized across the cell – responds linearly to the applied electric field. We find that, when measuring anisotropy, response and forgetting both have a single time constant *τ*_forget_. However, the directionality *D*(*t*) is a nonlinear function of *A*(*t*) – which means that the apparent dynamics of the directionality can be very different from the dynamics of the anisotropy. As a consequence, fitting *D*(*t*) can be potentially misleading. In Fig. 3, we show the “apparent” response and forgetting times from measuring the time *D* takes to reach its asymptotic values. We see, consistent with the results of [4], that the apparent response time decreases with electric field, and the apparent forgetting time increases with electric field, leading to these two values diverging. However, if we were measuring the underlying distribution of sensors instead, and we were in the weak-field limit of *κ <* 1, we would expect that the response time and forgetting time would be identical, as we previously noted.

**FIG. 3.**
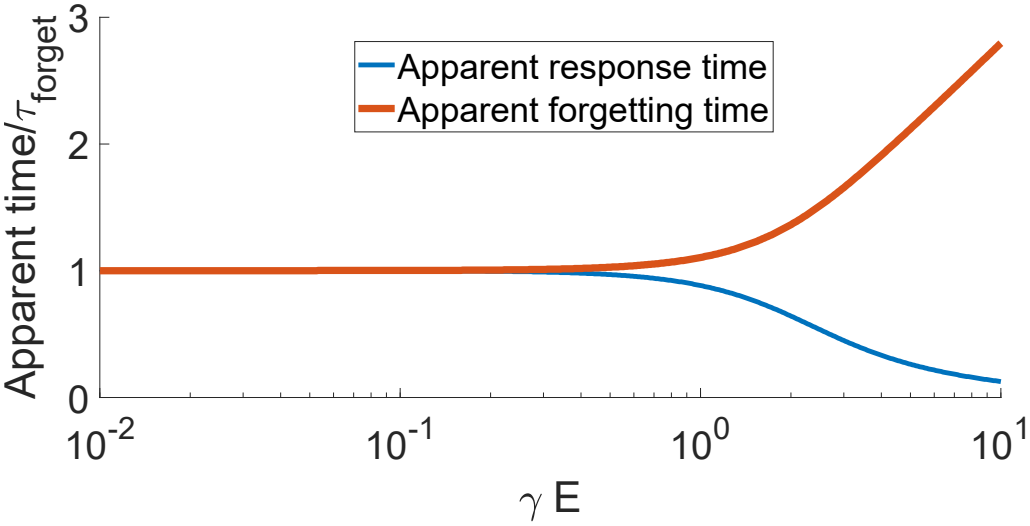
The apparent time for a cell to forget a signal or respond to an applied field can differ from *τ*_forget_ if it is computed by measuring the time for the directionality to respond. Curves show the apparent time from fitting to *D*(*t*) given by Eq. 9 for a field stepped up or down. This is solely a function of the rescaled electric field *γE*_0_ with 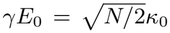. The apparent time to forget when measuring directionality is the time when *D*(*t*) reaches 1*/e* of its initial value. The apparent time to respond is the first time where *D* (*t*) reaches 1*−*1*/e* of its long-time value.

The intuitive reason for the decreased response time as *γE*_0_ is increased is because the strength of the electric field and the number of sensors change how sharply the directionality *D* varies when the field is turned on via Eq. 9. Large values of *E*_0_ lead to large values of *κ*_0_, which will lead to a quickly-saturating directionality as plotted in Figure 2b – directionality can quickly reach its maximum value before sensors become fully anisotropic. However, even if the electric field strength is relatively weak and the sensor anisotropy is small, if the number of sensors *N* is large, small changes in anisotropy will create a large change in directionality – again creating a short apparent response time. Similarly, if the field strength is very high or the number of sensors is very high, the anisotropy must decrease very far from its maximum value in order of the directionality to decrease away from its saturating value (see, e.g. Fig. 2d), increasing the forgetting time.

Previous work has argued that a fast (minute-scale) response of cells in galvanotaxis should be interpreted as evidence against the idea of a sensor being transported across the surface of the cell [9]. We see in Fig. 2 and Fig. 3 that fast response is not prohibited in a model with electromigration of sensors. In fact, the directionality of a cell can respond much faster than the timescale for diffusion of sensors across the entire cell. Essentially, this is because even a small change in sensor distribution still provides enough information for the cell to follow. This shows that it is in principle possible for a cell’s direction to respond much faster than *τ*_forget_ even if cells use only sensor electromigration, but it does not rule out potential other factors in the experiments of [9], such as the role of ion channels.

## V. FITTING RESPONSE FOR SENSORS AND DIRECTIONALITY SIMULTANEOUSLY CONSTRAINS MODEL

Our model describes the cell’s directionality in terms of the anisotropy of sensor positions *A*(*t*). If we can measure the sensor anisotropy *A*(*t*) and the cell’s directionality *D*(*t*) simultaneously, we can fit our model to determine the three parameters that will completely characterize the cell’s response – *N, κ*, and *τ*_forget_. We study the redistribution of the sensor galvanin (TMEM154) [10]. We re-analyze data collected by [10], who quantified the redistribution of galvanin tagged with a green fluorescent protein (GFP) in HL60 neutrophil-like cells exposed to electric fields, while also tracking the cell’s motion. We compute the ratio of cathodal to anodal fluorescence of GFP-galvanin (Fig. 4). When there is no field, the ratio of cathodal to anodal fluorescence is roughly unity, and the cell has *D*(*t*) ≈0, as expected. When the field turns on, galvanin redistributes to the anodal side of the cell, and the cathode-to-anode ratio decreases. Simultaneously, the cell’s directionality increases (Fig. 4). We fit the experimental cathode/anode ratio and *D*(*t*) data for *t <* 600 s to our model, assuming that the ratio Cathode/Anode 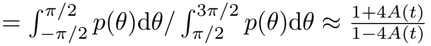, and then predict the cell’s behavior after the field is turned off (*t >* 600 s). We see, as expected from the theory, that the sensor anisotropy timescale to forget is exactly that of the sensor anisotropy’s time to respond. We also see excellent agreement between our prediction and the observed directionality after the field is turned off.

**FIG. 4.**
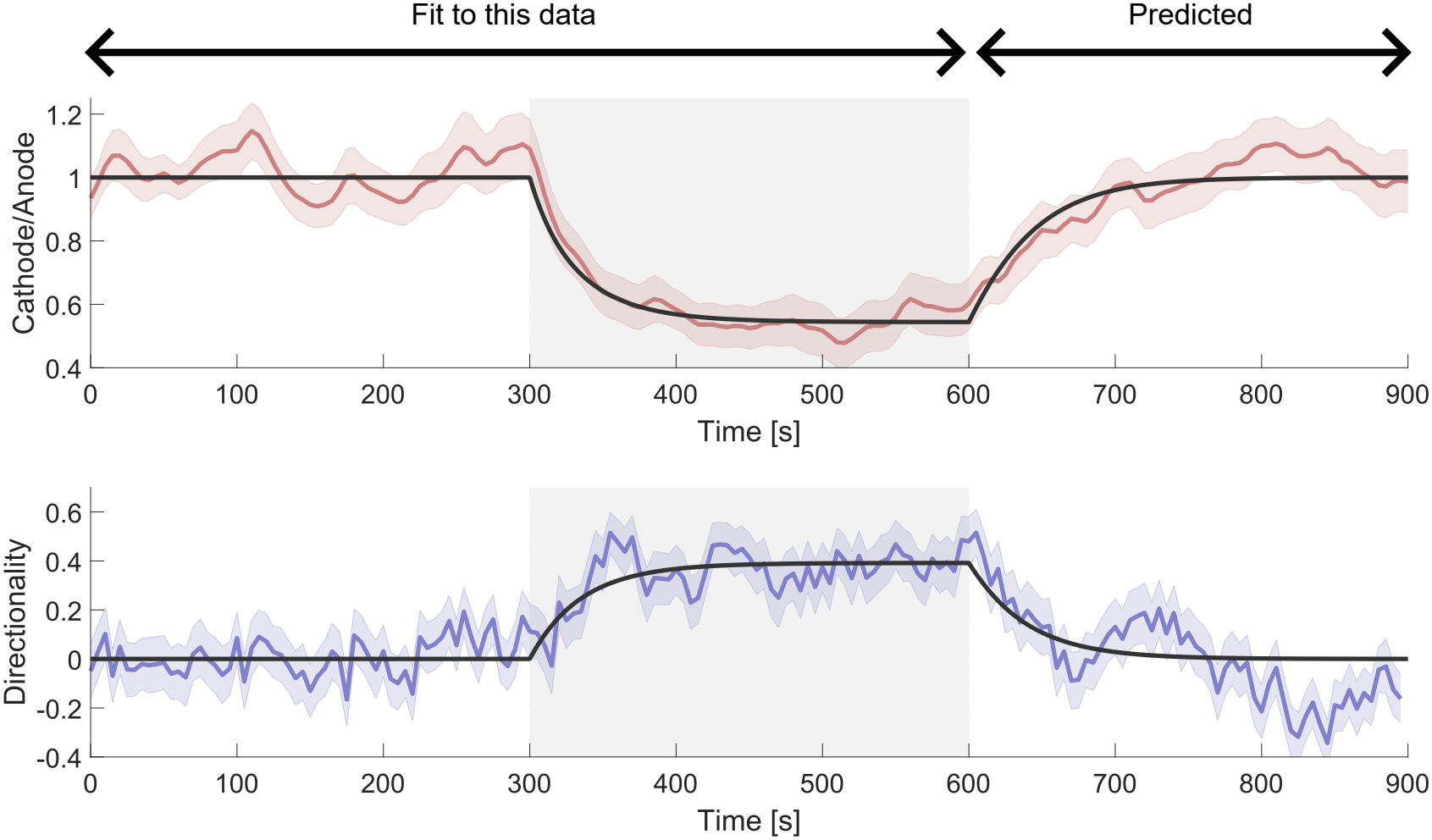
Theory is able to predict experimental results. Colored curves are experimental data originally taken in [10], lines are our theory. Anisotropy is quantified with the ratio of the average of cathodal fluorescence to anodal fluorescence (see Methods). We fit this to 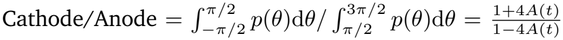 with *A*(*t*) given by Eq. 5, and fit the directionality to Eq. 9 with the appropriate *A*(*t*). *n* = 46 cells; error bars are the standard deviation of the bootstrap distribution (roughly a 68% confidence interval).

The parameters extracted from this fit are *κ* = −0.46 ±0.09, *τ*_forget_ = 37 ±10 seconds, and *N* = 4.8 ±3. We see that the value of *κ* is in the region where our linear theory agrees with stochastic simulations (Fig. 2), suggesting it is self-consistent to apply the linear theory. The timescale *τ*_forget_ is shorter than we would initially guess. The longest extent of the neutrophil is roughly ~20 µm, and the diffusion coefficient of galvanin has been measured to be ≈0.5 µm^2^*/*s in HL-60, so if we chose *R* = 10 µm, we’d expect *τ*_forget_ ≈200 seconds. We believe this discrepancy reflects the fact that HL-60 is not circular in shape. In fact, we can generalize our linear response results to an arbitrarily-shaped cell (Appendix C), and find that for a cell with characteristic length *L*, we’d expect dynamics similar to that of Eq. 4 but with a time scale

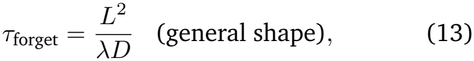

where *λ* is a geometric parameter – the smallest relevant nonzero eigenvalue of the Laplacian on the surface of the cell. For a spherical cell, this length scale is the cell radius *R*, and *λ* = 2, predicting *τ*_forget_ = 100 s. For a cell with maximum radius *L* = 10 microns but other dimensions (height, width) shorter, we find using finite element methods [40] still larger values of *λ*, since this decreases the time to diffuse across the cell. We estimate a relevant time to respond, depending on cell size and orientation, of 20− 80 seconds (Appendix C). We thus argue that the experimental value of *τ*_forget_ ≈37 s is consistent with our model, which includes only simple diffusion. Extensions of our model including anomalous or active diffusion, which have been observed for membrane-bound proteins [41, 42], but not immediately observed in galvanin by [10], could also potentially alter, e.g. the scaling of *τ*_forget_ with cell size. Testing these assumptions more carefully is a potential future area of work. We also note that the experimental value of the diffusion coefficient of galvanin *D*≈ 0.5 µm^2^*/*s was quantified in immobilized (latrunculin-treated) cells – in motile cells, sensor-cortex interaction or redistribution of the sensor by cortical flow [43] may also potentially be relevant.

The fit value of the number of galvanin *N*≈ 5 is obviously not the true number of galvanin on the surface. If *N* were this small, we would be able to resolve individual galvanin as diffraction-limited spots. This result strongly argues that the limiting factor of electric field sensing in HL-60 is *not* the finite number of galvanin, as we had previously assumed in [32, 33]. Instead, we must think of *N* as an effective number *N*_eff_ as we discussed above, where the effective number of sensors is really set by additional sources of noise. An initial guess for this value of *N*_eff_ is that this reflects a typical value of the number of random protrusions present on a HL-60, with these random protrusions uncorrelated with a field. More detailed models to fit this parameter would attempt to resolve stochasticity in cell polarity [44–46], with the polarity coupled to sensor redistribution (as, e.g. by [47]).

A potential caveat to our fitting in Fig. 4 is that we have assumed *D*_max_ = 1, i.e. that directionality will only saturate once it reaches unity. Decreasing *D*_max_ will increase the value of *N* in our fit (Appendix D) – either decreasing *N* or decreasing *D*_max_ in Eq. 12 will reduce directionality, so there is some confounding between these two parameters. However, unless the maximum directionality is by coincidence nearly exactly the largest directionality we observe – which is in conflict with other measurements – this effect will not change the core conclusion that noise from finite sensor number is likely not limiting (Appendix D). We also discuss some potential other ways of fitting this curve in Appendix D, such as using our results for a spherical cell, or what happens if we define the Cathode/Anode ratio to exclude some portion of the cell boundary, as done in the experiments (see Methods). In general, we find different geometric assumptions can change the particular values of *κ* for a fit but do not change the scale of the response time or the order of magnitude of *N*_eff_, so we do not think these will affect our core conclusions.

We note in Fig. 4 that the apparent response time and forgetting time for the directionality are quite similar, as found at relatively low field strength in Fig. 3. This behavior is expected since the time to respond and time to forget begin to differ first when directionality nears its saturation value, when |*γE*| ≳ 1. For the fit parameters for the HL-60 experiment, we find |*γE*| to be 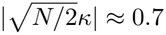.

## VI. RESPONSE TIMES DEPEND ONLY WEAKLY ON MEDIUM VISCOSITY

One key piece of evidence that suggests sensors have to redistribute around the cell’s surface for a cell to respond to an electric field is the observation of Allen et al., who found that increasing the viscosity of their keratocyte’s solution from 1 cP to 50 cP increased the time to respond [4]. We note that these earlier experiments are on a different cell type – fish keratocytes – and also have a significantly longer response time, of order of ten minutes. We replot the response curves for keratocytes in 1 cP and 50 cP solutions from [4] in Fig. 5a, and fit the curves to our model (dashed lines). Surprisingly, even though the viscosity of the medium is increased by a factor of 50, the change in response time is relatively moderate and clearly not linearly proportional to the change in viscosity. The free parameters of our model are *N, κ*_0_, and *τ*_forget_; because we only have experimental measurements of directionality for keratocytes, which depends on anisotropy as 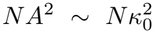, we cannot separately fit *κ*_0_ and *N* and can only fit 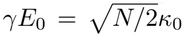. However, we can still use this model to extract the timescale *τ*_forget_, finding *τ*_forget_ = 7.3 min for 1 cP and 16.2 min for 50 cP. Our model says that *τ*_forget_ = *R*^2^*/D* – so we can explicitly determine the ratio of diffusion coefficients of the sensor at these two solution viscosities, 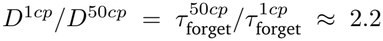. Is this consistent with the known effect of extracellular viscosity on the diffusion of membrane proteins? The diffusion of membrane proteins is commonly described with the Saffman-Delbruck-Hughes-Pailthorpe-White (SDHPW) model [48–51], which takes the form

**FIG. 5.**
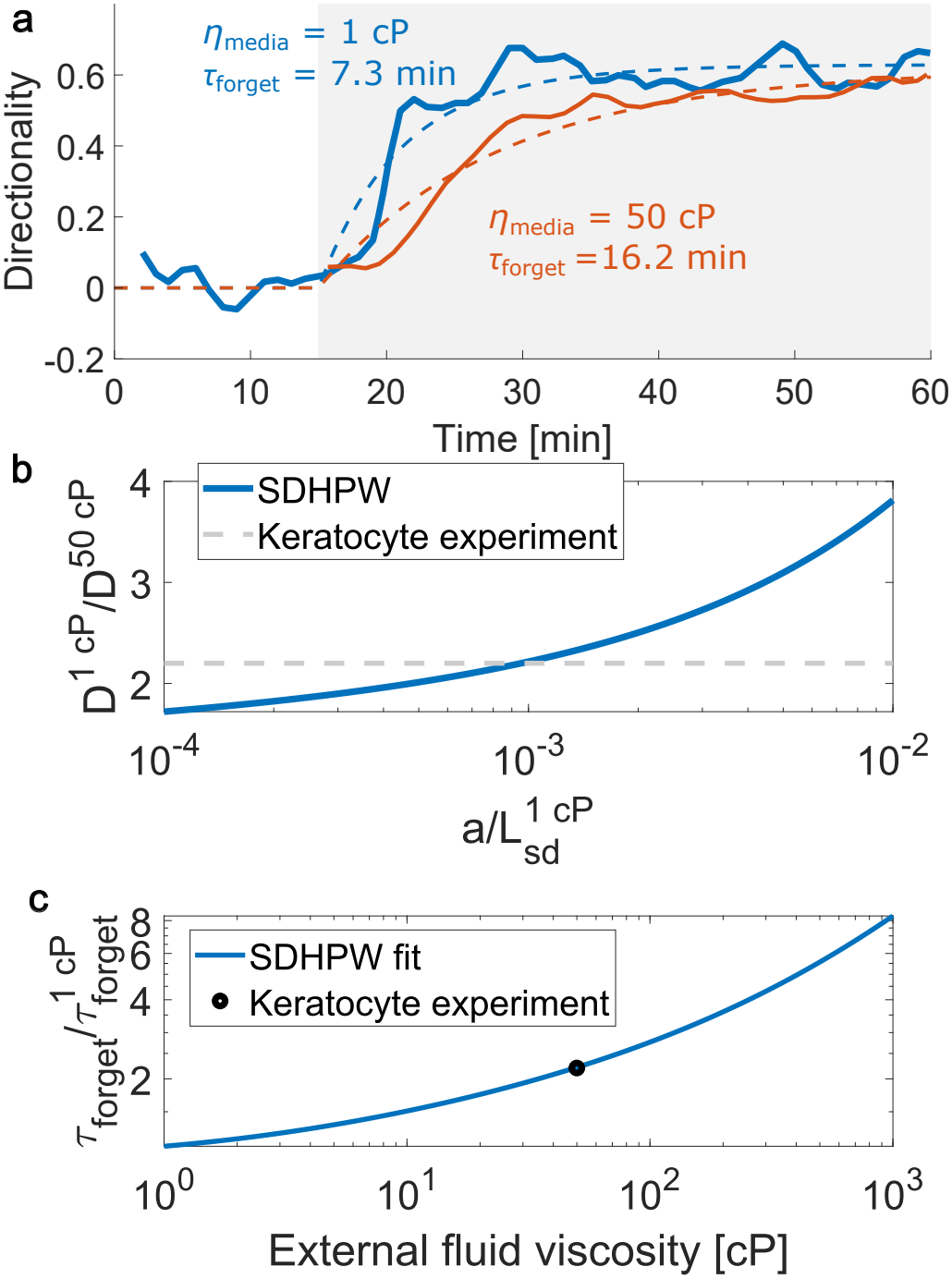
a) Directionality as a function of time for experiments on keratocytes in normal media (viscosity = 1 cP) and high-viscosity media (50 cP) when an electric field is turned on at *t*_on_ = 15 min. Solid lines are data from [4]; dashed lines are fits to Eq. 9 with Eq. 5. b) Dependence of the ratio of sensor diffusion coefficients as a function of the unknown parameter 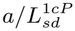 from the SDHPW theory, Eq. 14 (using the full form of [48]), with the value of 2.2 implied by the keratocyte experiments as a dashed line. c) Predicted time to forget relative to the normal-viscosity time as external fluid viscosity is varied, given Eq. 14, and assuming 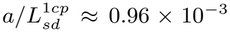 found in panel b).

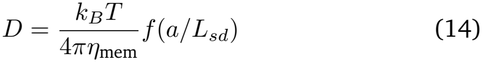

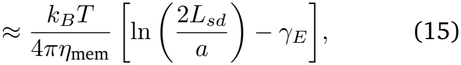

where *L*_*sd*_ = *η*_mem_*/*(*η*_*f*_ + *η*_cyt_) is the Saffman-Delbrück length, *η*_mem_ is the membrane’s surface viscosity, *η*_*f*_ the solution viscosity, *η*_cyt_ the cytosolic viscosity and *a* the radius of the protein, and *f* is a complicated function [48, 49, 51]; *γ*_*E*_≈ 0.577 is the Euler-Mascheroni constant. In the limit of small *x, f* (*x*) is only logarithmically dependent on *x* (Eq. 15), so we expect a relatively weak dependence on fluid viscosity.

The experiments imply that the ratio *D*^1*cp*^*/D*^50*cp*^ ≈ 2.2 – is this consistent with the SDHPW model in Eq. 14? The prediction for this ratio depends on an unknown unitless parameter 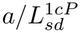, the ratio of sensor protein radius *a* to the Saffman-Delbrück length for a membrane in water. We can get a rough estimate of plausible ranges for this value. We expect protein sizes *a*~ nm, and the Saffman-Delbruck length *L*_*sd*_ to be on the order of microns if *η*_*f*_ has the viscosity of water [52], so the most likely value is of the order 10^*−*3^ – though this is a loose estimate. We show in Fig. 5b that the diffusion ratio is reasonably close to the experimental value of 2.2 for a broad range of values of the unknown 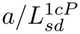. We then pick the value of 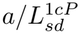 that produces the experimental value of 2.2 (the intersection of the two lines in Fig. 5b), finding 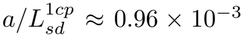. Given this fit parameter, we predict the dependence of the timescale *τ*_forget_ on the external fluid viscosity *η*_*f*_ (Fig. 5c). We see a relatively weak, logarithmic dependence, where even an increase of three orders of magnitude in medium viscosity would only increase the response time by a factor of 8.

Using the SDHPW model for diffusion in a cell membrane may be a little simplistic – the SDHPW model describes the diffusion of a cylindrical inclusion in a two-dimensional homogeneous membrane. This model neglects the role of, e.g. protein inclusions [53, 54], coupling to the cell cortex [55], the extracellular part of the protein, finite size of the cell [56], potential viscoelasticity of the membrane [57], and heterogeneity of the membrane. These generalizations would change the particular form of the curves in Fig. 5bc. However, many of these generalizations will also have a similarly weak dependence on extracellular viscosity in many limits – the core reason for this is that because membrane viscosity is so high relative to the viscosity of the outside fluid that the drag from the outside fluid is overwhelmed relative to the drag due to the membrane. We have also assumed that the outside fluid can be well-described by a continuum fluid with a fixed viscosity – this is not necessarily the case in galvanotaxis, as discovered by [6], who found that polymer solutions with the same viscosity but different polymer structure can alter galvanotaxis in different ways.

If keratocytes also use galvanin as a sensor (as supported by the reduced galvanotaxis in galvanin knockout keratocytes in [10]), we would expect that *τ*_forget_ would also be on the order of *R*^2^*/D* ~ 200 s since their typical spread area of ~400 µm^2^ [58] would correspond to an equivalent circle of radius ~11 microns. As we address in Appendix C, this is sensitive to the full three-dimensional cell shape – but it is still surprising that keratocytes have a slower response than HL-60 if protein diffusion limits both. There are two clear possibilities. First, it is possible that the sensor in keratocytes is not galvanin, or that galvanin has a slower diffusion coefficient due to, e.g. different interactions with other membrane proteins. These changes could easily lead to quantitative differences in response time. Second, it is possible that keratocytes take a long time to repolarize relative to the sensor redistribution, and it is this time that is setting the seven-minute time scale. This would be in conflict with our model and could be challenged by experiments testing whether, e.g. keratocytes obey our proposed limits for responding to pulsed electric fields. However, if this second option is true, there would need to be an explanation for the role of the external fluid viscosity in setting this timescale – possibly in the nontrivial effects of viscosity found by e.g. [59, 60].

## VII. PULSED FIELDS

While applied electrical fields may help guide cells in migration toward the wound and promote wound healing [1, 61], applied fields also have the risk of damaging cells due to local disruption of pH due to electrochemical effects [62, 63]. As a result, there is a clear rationale for applying the weakest sufficient field and managing exposure time. One route of doing this is applying a pulsed DC field instead of a constant DC field [63]. Experiments by Ren et al. found that pulsed electrical fields can still effectively guide keratinocytes, though there is a dose-response relationship between the duty cycle of the pulsing and the directionality of the cells [63]. Ren at al. also observed that changing the frequency of the pulsing over a broad range (0.005 Hz - 5000 Hz) did not create a large change in the directionality of the cells. Can we understand these results in our framework?

We show the results of a simulation of a cell exposed to a pulsed field in Fig. 6. We assume that the field is periodic, with a duty cycle DC, and a period *T*

**FIG. 6.**
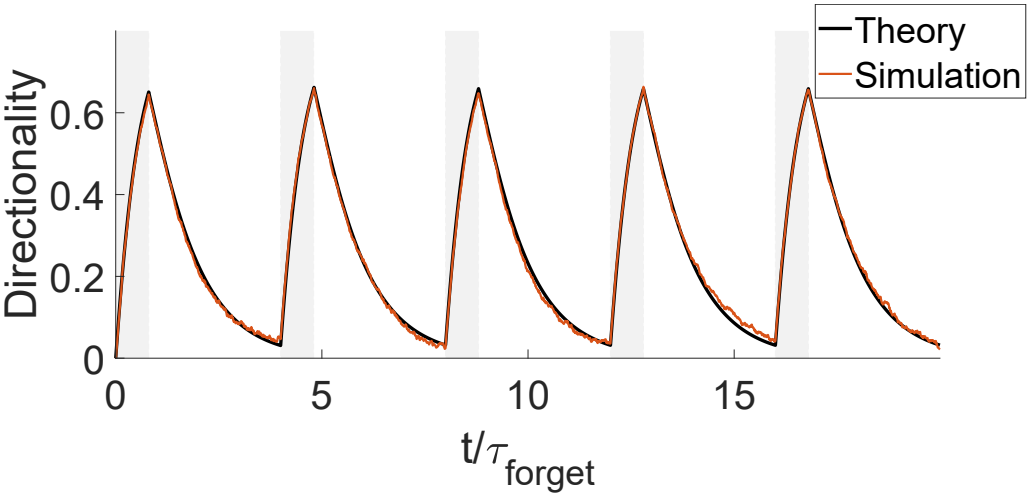
Illustrative example of the average of a cell’s directionality in a pulsed field. Shaded region indicates when the field is on. Period *T* = 4*τ*_forget_, duty cycle = 20%, *κ*_0_ = 0.1, *N* = 1000. Directionality is averaged over 5000 simulated cells. Simulation is the stochastic sensor simulation as in Fig. 2.

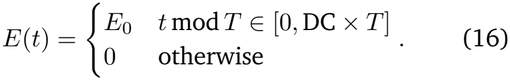

We see, as we expect, that the directionality increases quickly while the field is applied, and then decays when the field is turned off. We see that the directionality also rapidly reaches a steady-state periodic form. We also see that our stochastic simulations and our simple theory agree well in the small-*κ*_0_ limit.

### A. Linear response theory predicts effective field is time-average of applied field

How can we describe the dynamics of a cell exposed to a pulsed field? Can we describe it as moving in some effective field strength that depends on the duty cycle and period of the pulsing? Our first natural guess would be that *E*_eff_ = DC ×*E*_0_ – a field at 50% duty cycle is like applying half the field (as, e.g. [5], have proposed). We can show that this guess is reasonable.

If we integrate the equation of motion for the anisotropy (Eq. 4) over one period, from *t*_*i*_ to *t*_*i*_ + *T*, we find

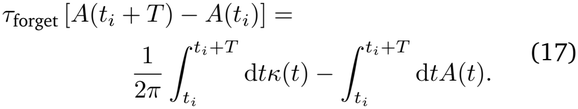

After an initial transient, the anisotropy will have period *T, A*(*t*_*i*_ + *T*) = *A*(*t*_*i*_) and the left-hand-side of this equation is zero. If we divide Eq. 17 by *T* we find that the time average over one period 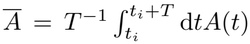 is just driven by the time-average of the electric field,

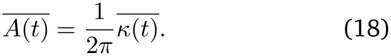

For a pulsed field with duty cycle DC, the time-averaged electric field is just *Ē* = *E*_eff_ = DC ×*E*_0_. We see then that the time-averaged anisotropy *Ā* is just proportional to DC × *E*_0_ – i.e., that on average, anisotropy is proportional to the effective field strength we defined earlier. However, this does not necessarily mean that the *directionality* is solely a function of the effective field strength.

### B. Directionality is limited by the effective field

One natural hypothesis, based on our earlier results on the anisotropy, is that we could assume that a pulsed field with duty cycle DC has directionality *D*(*Ē*). We test this hypothesis in Fig. 7. We plot time-averaged directionalities computed from the full time-dependent dynamics of Eq. 9 for different values of field strength *κ* and duty cycle DC and plot them as a function of *Ē* = DC × *E*_0_. If time-averaged directionality *D̄* were solely a function of *E*, we’d expect these results to all collapse onto a single curve, given by Eq. 8 with a constant *Ē* (dashed line). Instead, we find that while many simulations collapse onto this line, larger values of *κ* lead to a time-averaged directionality lower than expected for a constant field of *Ē*. We emphasize that this is not because of a breakdown of our small-*κ* approximation – all of the directionality curves are computed by the linear response theory and are in the limit where theory and stochastic simulations agree well.

**FIG. 7.**
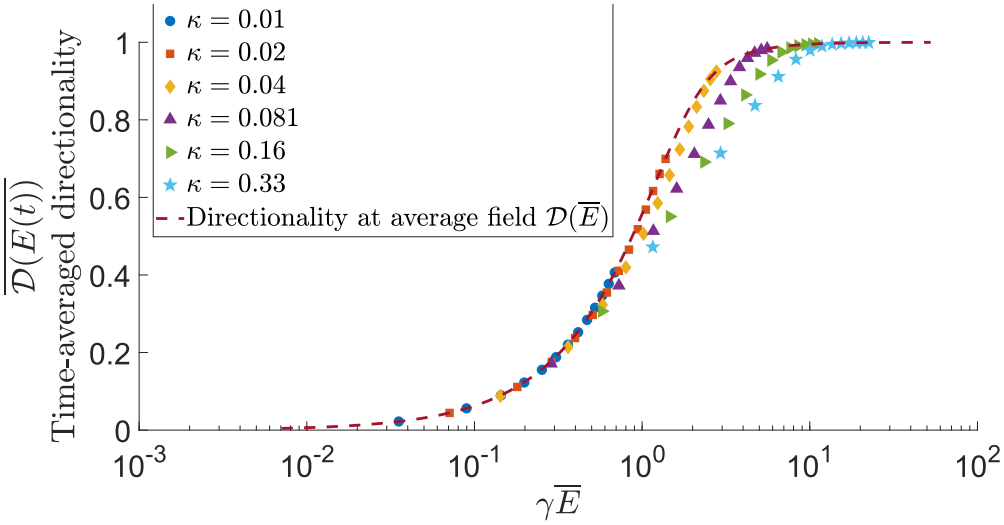
Time-averaged directionality 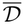 is bounded by *D*(*E*_eff_) where *E*_eff_ = *Ē* = DC*× E*_0_. These results are computed from numerically evaluating Eq. 9 with a pulsed field with varying DC (ranging from 5%-100%) and varying *κ*. Period of pulsing is *T* = 4*τ*_forget_; simulation is over total time of 40 *τ*_forget_, and the time-averaged directionality measured over the last two periods of the simulation to avoid initial condition effects. 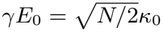.

In Fig. 7, we see that the time-averaged directionality is less than or equal to the directionality predicted at the time-averaged field *E*_eff_,

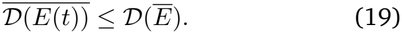

Eq. 19 means that there is no free lunch – if you pulse the field to be on x% of the time, you will be no more directional than the same field at x% strength. In fact, Eq. 19 is always true for any periodic field where *A*(*t*) does not change signs, and not just for a pulsed field – Eq. 19 is simply Jensen’s inequality applied to the concave function *D*(*E*) (Appendix F).

We see from Fig. 7 and Eq. 19 that *D*(*Ē*) is always a limit to galvanotaxis in a pulsed field – but in many cases cells are nearing this limit. We argue that there are two possible reasons why *D*(*E*_eff_) can be a good approximation to the time-averaged directionality: First, if the electric field is sufficiently weak so *D* does not reach its saturating value of 1, can be approximated as nearly linear in the anisotropy *A*(*t*). If *D*~*A*(*t*), then the time-averaged directionality will naturally be proportional to the time-averaged anisotropy, which is proportional to the time-averaged electric field. However, we see that in Fig. 7 that we can still have collapse to the effective curve even in cases where the directionality is near unity. We think this happens if the duty cycle is close enough to 1 that the directionality as a function of time is relatively constant. In that case, the directionality should be close to the directionality given the time-averaged anisotropy.

We study the phase diagram of when the time-averaged directionality agrees with the directionality at the effective field more carefully in Fig. 8. We see that largely, the time-averaged directionality is well-approximated by the directionality at the effective field strength. However, deviations occur when the duty cycle is small and *κ* relatively large (~0.1 − 1). This is exactly what we would expect – this is the range where the field strength is large enough so that the directionality occasionally saturates, but that the time between pulses is long enough relative to *τ*_forget_ for the directionality to decay (Fig. 8, point A). If the pulsing has a higher duty cycle (point B), directionality is roughly constant. At lower *κ*, directionality is better-approximated as linear in field strength, and even though directionality is strongly time-varying, its time average is close to *D*(*E*_eff_) (points C, D).

**FIG. 8.**
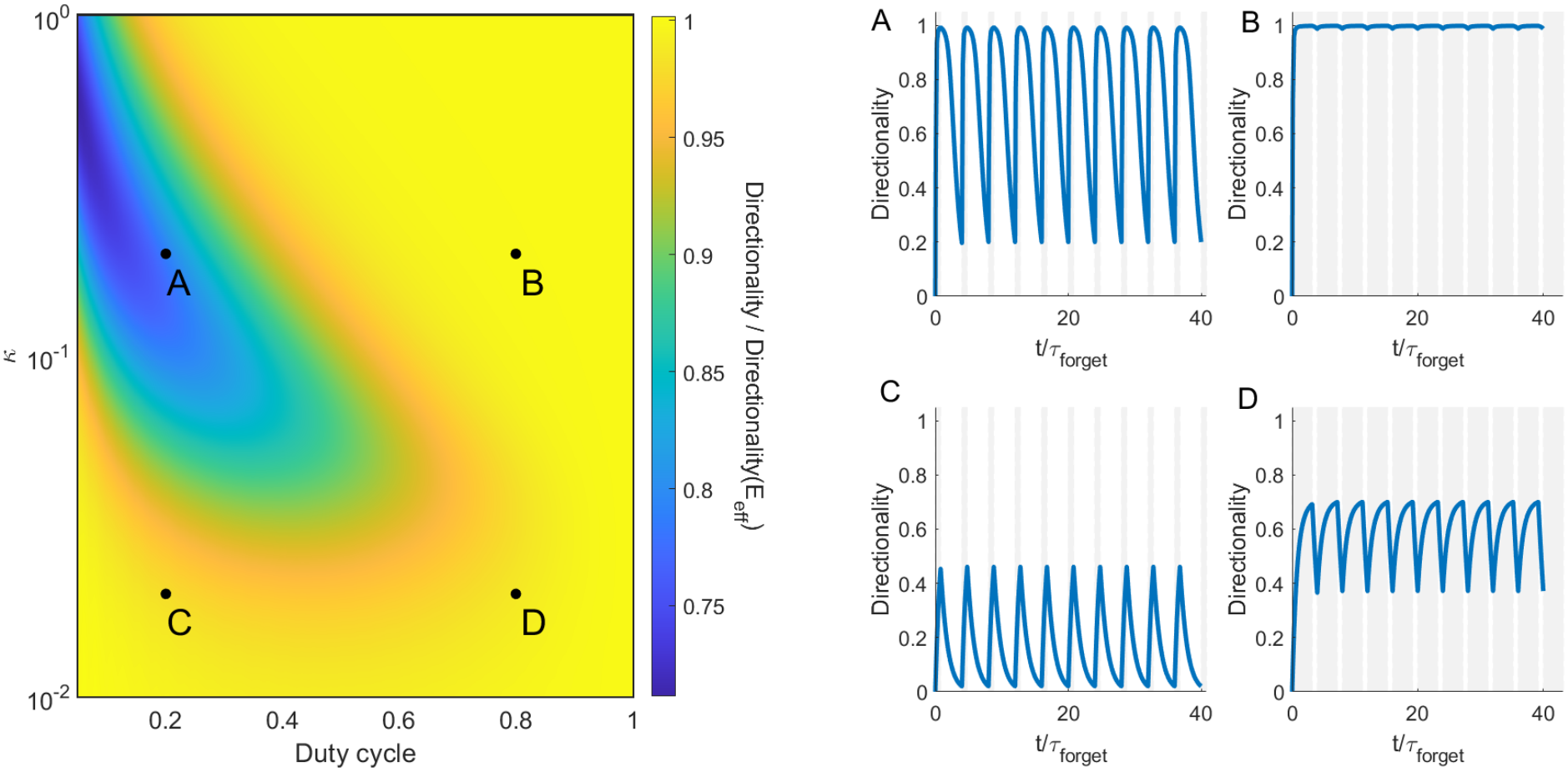
Phase diagram showing where time-averaged directionality 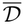 can be lower than the directionality at the time-averaged field, *D*(*Ē*). Points *A, B, C*, and *D* are illustrated to the right, showing instantaneous directionality as a function of time. Shaded regions show when the field is on. These are computed from Eq. 9. Period *T* = 4*τ*_forget_, total run time is 40*τ*_forget_, Δ*t* = 10^*−*3^*τ*_forget_, and the time-averaged directionality is from averaging over the last two periods.

We note that this deviation requires that the period *T* is larger than *τ*_forget_. If the pulsing period is much less than the forgetting time, the directionality is nearly constant and well-approximated by the directionality at the effective field (Fig. S2).

### C. Experiments on pulsed field are consistent with cells responding to a time-averaged field strength

Ren et al. [63] exposed keratinocytes to pulsed fields with different field strengths and duty cycles, and measured their directionality. These experiments have frequency *f* = 0.1 Hz (*T* = 10 s), so we expect them to be in the limit of *T* ≪*τ*_forget_, and expect 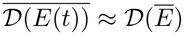. We replot the measurements of [63] in terms of directionality as a function of the effective field *E*_eff_ = DC ×*E*_0_ in Fig. 9. We see that the experimental data does collapse – though roughly – onto a single curve, suggesting that our theory is reasonable for this case. However, we do find that no matter the electric field strength, the keratinocyte’s directionality does not come close to one. This can happen, as we mentioned in Section III A, if a cell even with perfect information about the field’s orientation does not follow the field perfectly, e.g. due to oscillations around the target direction or irreducible noise in the motility process. The simplest way to model this is to assume that the cell travels in a direction that has some random error from its best estimate. This has the effect of reducing the maximum directionality [32], so instead of Eq. 9, we introduce a maximum directionality *D*_max_, following Eq. 12, and fit both *γ* and *D*_max_ to the data in Fig. 9. We find *γ* = 0.030 mm/mV, and *D*_max_ = 0.68 (dashed line in Fig. 9).

**FIG. 9.**
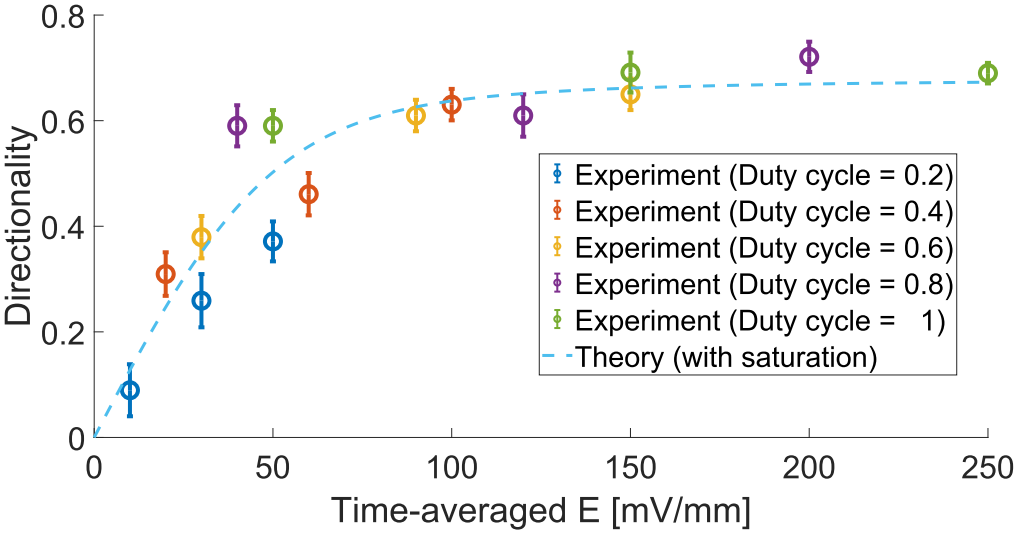
Directionality data at different duty cycles and electric fields, at a frequency of 0.1 Hz, from Fig. 4A of [63] roughly collapse as a function of the time-averaged *E, Ē* = *E ×* DC.

This collapse and fit is relatively rough by the scale of the error bars. This could reflect additional variability not necessarily well-summarized by the standard error bar, e.g. day-to-day variability in motility experiments [64], or could reflect that there are additional effects of pulsed electric field not captured in our model. We have also found in our simulations that it is easy to have systematic errors arising from not averaging over a complete period of a pulsed signal, which is another possible reason for this deviation, since the time scale over which directionality is measured is not clear in [63]. Additional experiments that resolve the directionality as a function of time, as in Fig. 4 and Fig. 6, would provide a stronger test of our model.

### D. Directionality is weakly dependent on period of pulsed field

Ren et al. [63] found that directionality is not significantly dependent on the frequency of the applied pulse, at least for frequencies in the range 5000-0.005 Hz. We find in our simulation that directionality is generally independent of period – as long as the period is shorter than *τ*_forget_ (Fig. 10). We identified in the previous section and in Fig. 8 conditions where the time-averaged directionality can be approximated as 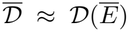. In these cases, we would expect time-averaged directionality to be independent of period since the time-averaged field strength is DC× *E* no matter what *T* is. However, as *T* gets larger and *T/τ*_forget_ becomes large, we expect 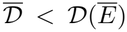, so we expect the time-averaged directionality 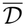 to decrease, as we see in Fig. 10. Ren’s results finding relatively weak dependence on frequency suggest that their experiments are in the range *T/τ*_forget_ *<* 1 over their entire frequency range, which argues that the *τ*_forget_ for keratinocytes is larger than 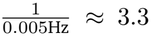 minutes. This timescale seems plausible given values of *τ*_forget_ we have seen in other systems, e.g. the keratocyte *τ*_forget_ of 7.3 minutes, but will of course depend on the specific diffusion coefficient of the sensor in keratinocytes and the geometry of the cell.

**FIG. 10.**
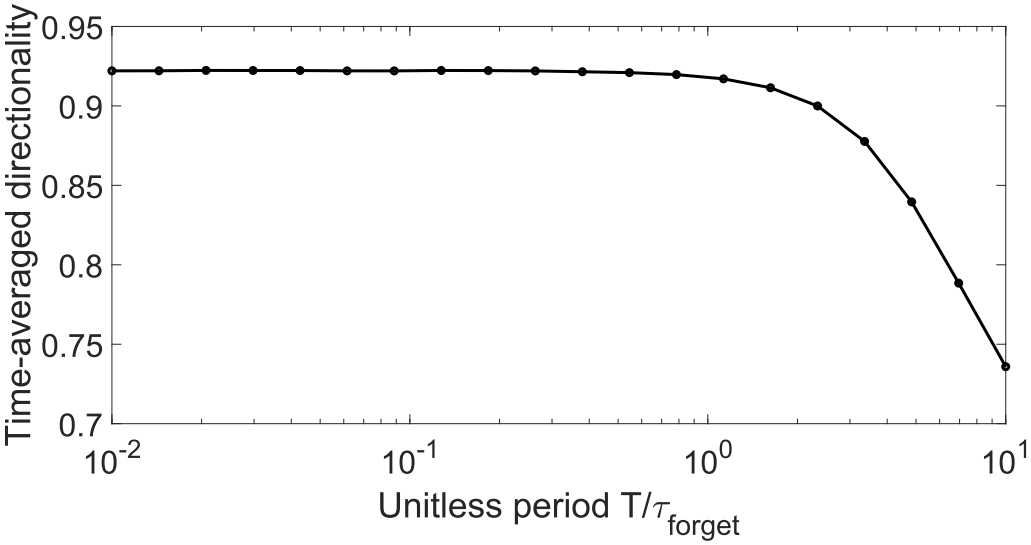
Time-averaged directionality 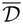 for a pulsed field as a function of the period *T*. Computed from Eq. 9 with *γ* = 0.030mm/mV (the value fit from Fig. 9), *E* = 150 mV/mm, and a 60% duty cycle. We assume *D*_max_ = 1 here; changing the value of *D*_max_ would simply scale the curve.

## VIII. DISCUSSION

Galvanotaxing cells are guided by the distribution of charged sensors on the surface of the cell. We show that the anisotropy of these sensors obeys a simple linear response relationship with the applied electric field *E*(*t*), with a timescale of response ~*R*^2^*/D*. However, the directionality of the cell’s migration is a *nonlinear* function of the anisotropy. Fitting this model to experimental data of [10], we immediately find that the effective number of sensors *N*_eff_ is below any plausible value, suggesting that galvanotaxis is not limited by intrinsic stochastic noise. We also see that we can predict the behavior of a cell when the field is turned off solely by its response dynamics to a switched-on field. Our results solve a large number of mysteries, including showing how a cell’s directionality can respond very quickly even though sensors must redistribute over the cell’s surface, and why a keratocyte’s response time is so weakly dependent on the environmental viscosity. In addition, our model shows there are strong constraints on the ability of a pulsed DC field to help guide cells – a field turned on 10% of the time will have no more effect on directionality than a field at 10% of the magnitude, which we find is consistent with experimental measurements.

By simultaneously quantifying cell directionality and galvanin redistribution, we argue that that galvanotaxis in HL-60 neutrophil-like cells at least is likely not limited by finite galvanin number. This analysis is based on the fits to our model, and could certainly be supported by additional experimental measurements, such as a rigorous quantification of and modification of the number of galvanin, which we hope to pursue in future work. The idea that shot noise from finite galvanin count is not relevant to setting directionality is also supported by the observation in [10] that galvanotaxis of wild-type HL-60 is quantitatively consistent with the galvanotaxis of galvanin-knockout HL-60 with overexpressed galvanin-GFP.

What is the source of noise represented by our parameter *N*_eff_ if it is not from finite galvanin number? We note that even in the absence of an applied electrical field, HL60 cells develop protrusions stochastically that can reorient the cell [10, 65]. Our expectation is that it is either the randomness of stochastic protrusions or randomness of cell polarity (see, e.g. [66, 67]) that can degrade galvanotactic directionality. Extensions of our model to address stochastic protrusion or polarity dynamics [45, 68–71] could address this possibility, at the cost of significantly complicating the model. For a model like this, the noise strengths would have to be calibrated to explain our observation of *N* ≈ 5. Another possibility would be that galvanin does not act as individual sensors, but as large aggregated units, potentially connected by lipid rafts as suggested by [72]. If this is the case, *N* ≈ 5 may reflect the number of independent sensing units.

It is somewhat surprising how well this simple model works to capture the dynamics of HL-60 and keratinocytes. Our assumption is that the sensor distribution instantaneously sets the cell’s direction – neglecting the complex dynamics of cell polarity [73, 74]. This excludes any memory that the cell could have. For instance, in many cells exposed to a chemotactic stimulus – including HL-60 – there is a significant (minutes) memory of a polarizing stimulus [75–77]. A memory of this sort would suggest that cells once polarized would, even once the field is turned off, continue with their original directionality with a slow decay. This is in contrast with the experimental results in Fig. 4. There are two clear possible explanations for this. First, it is possible that galvanin redistribution by an electric field is a “stronger” stimulus than a chemotactic gradient – i.e. it more effectively overrides the cell’s polarity. This is certainly consistent with the idea that galvanotaxis can be an overriding signal [2]. However, there may be a more prosaic explanation as well – the experiments of [76] that identified a memory in HL-60 were conducted in microchannel-based confinement. Cells in confinement are often observed to be more persistent [78], allowing them to override conflicting chemical cues [79, 80]. If we included an explicit memory or cell polarity within our model, we would expect to see response to pulsed fields with slower-decaying directionality – allowing cells to maintain a persistent directionality larger than *D*(*E*_eff_). We think that our results have created a reasonable “null model” for galvanotaxis – deviations from our constraints indicate interesting cell memory features.

Our work is solely focused on single-cell responses to electric field. If collective galvanotaxis arises from each cell individually responding to the field [81, 82], we would expect the collective dynamics to reflect single-cell dynamics, unless there were large degrees of cell-cell interaction that can slow or challenge external control [82, 83]. However, there are initial hints that collective dynamics may be qualitatively different from a collection of single-cell sensors. For instance, epithelial MDCK cells do not galvanotax in isolation, though groups can galvanotax [84]; similar results are found in neural crest cells [85]. In neural crest cells, collective galvanotaxis depends on a voltage-sensitive phosphatase [85], suggesting a substantially different mechanism than we have proposed here for single cells. In this case, collective galvanotaxis might have to be understood with theories describing how collective response can emerge from groups of cells that cannot individually galvanotax [86, 87].

## IX. METHODS

### A. Stochastic simulations

Symbols in Fig. 2 are stochastic simulations of sensor dynamics. An equivalent way to describe sensor motion to the Smoluchowski-like equation of Eq. 1 is to think of *N* independent sensors, with each sensor’s location *θ*_*i*_ obeying Brownian dynamics,

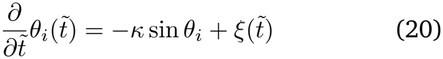

where the first term represents the sensor’s electromigration, which will take the sensor toward 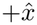 if *κ >* 0 and 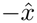 if *κ <* 0 and the second term controls the diffusion of the sensor. We also define 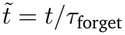 and 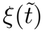 is a Gaussian white noise 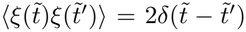. We follow our approach of [32], evolving the Brownian dynamics equation 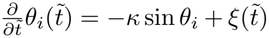 using the Euler-Maruyama method,

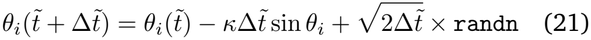

where randn indicates a random Gaussian number with mean zero and unit variance. 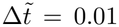 unless stated elsewhere. We evolve our *N* sensors independently. We compute the sensor anisotropy *A*(*t*) by first computing *p*(*θ, t*) with a histogram, and then computing 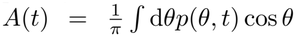. (We could also compute 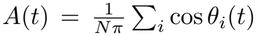 and find the same results.) We compute the directionality *D* by computing a vector in the direction of the vector sum, *ρ* = (*C, S*) with *C* = ∑_*i*_ cos *θ*_*i*_, *S* = ∑_*i*_ sin *θ*_*i*_, and then computing the ensemble average of 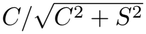 over *N*_its_ = 5000 iterations of the simulation.

### B. Experiments

Experiments were originally reported in [10]. Briefly, HL-60 neutrophil-like cells expressing galvanin-GFP were confined to a single plane on a glass coverslip using an agarose overlay. The cells were exposed to an electric field with a strength of 300 mV/mm, and fluorescence microscopy was used to monitor the redistribution of galvanin. Fluorescence measurements from individual cells were corrected for background fluorescence originating from the imaging setup and normalized to their maximum fluorescence intensity. The normalized fluorescence intensities at the anode and cathode (defined as in [10], where the cathode fluorescence is averaged over segments from −60 to 60 degrees, and the anode over 120 to 240 degrees) were averaged across individual cells and used to calculate the cathode-to-anode fluorescence ratio. Directionality was calculated from cell centroid trajectories using a moving average window of 8 frames, then computing the velocity from the averaged centroid trajectories, and the directionality from that velocity. Images were acquired every 5 seconds over a 15-minute period, during which cells were exposed to the electric field (300 mV/mm) between 5 minutes and 10 minutes.

## ACKNOWLEDGMENTS

We acknowledge support from NSF PHY 2412941, NIH R35 GM142847, and NIH K99GM147355. J.A.T. is supported by the Howard Hughes Medical Institute. N.M. acknowledges support from the Rowland Summer Research Fellowship. We thank Pedrom Zadeh, Thomas Brew, and Cailing Wang for useful suggestions and discussions, Wei Wang for a close reading of the paper, and Alan Lindsay for pointing out Ref. [88].

## Appendix A: Variances of estimated direction

In our default model, we assume that cells travel in a direction *ρ* = ∑_*i*_ (cos *θ*_*i*_, sin *θ*_*i*_) = (*C, S*), with *C* = ∑_*i*_ cos *θ*_*i*_ and *S* = ∑_*i*_ sin *θ*_*i*_. (This assumes *μ* > 0, i.e. the sensors go to the cell front; if the sensors travel to the cell back, *ρ* = ™(*C, S*); see also Appendix B.) We can compute the averages and standard deviations of *C* and *S*, assuming that 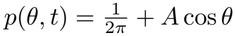, which are:

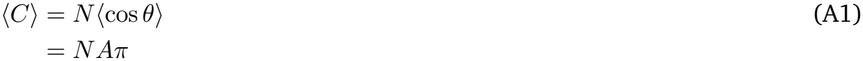

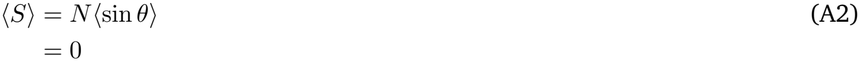

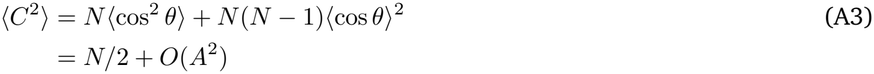

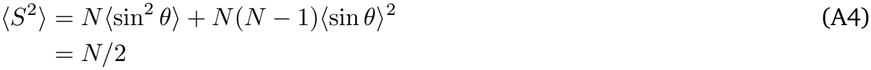

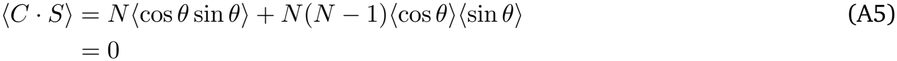

If we instead assume cells are traveling in a direction

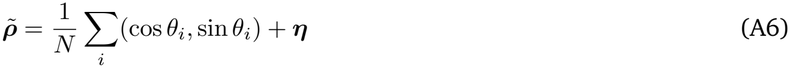

with ***η*** a vector of two independent Gaussian noises, both independent of the sensor locations then

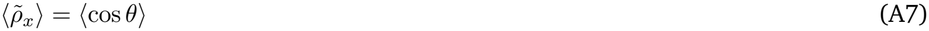

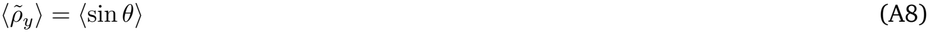

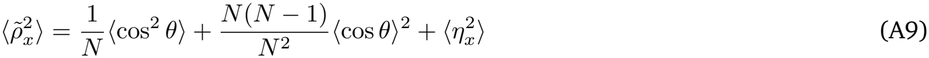

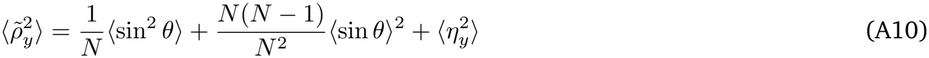

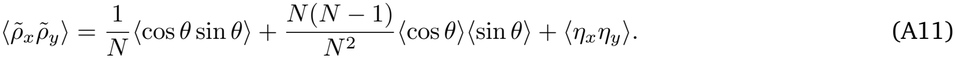

Assuming 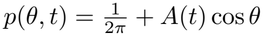, we find

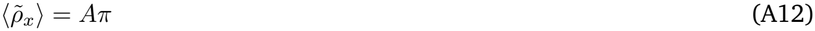

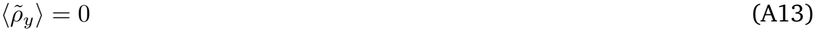

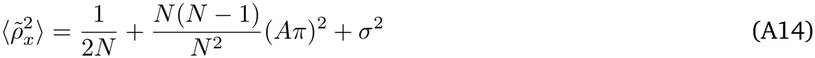

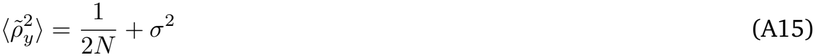

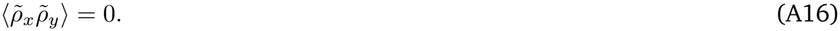

As we did in the main text, we neglect the terms of order *A*^2^. In this case, we can see that 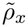 and 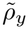 both have a variance of ^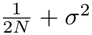^. We note that this limit of neglecting the term proportional to *A*^2^ will not be correct if we have *N* → ∞ and *σ* → 0 – at small enough noise, the difference between 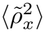 and 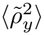 may matter, in which case the Bessel function form for *D*(*A*) we use in the main text (Eq. 9) would have to be generalized.

## Appendix B: Derivation of the directionality formula

Eq. 8 in the main text was originally presented by [36]. For completeness, we include a brief derivation of it here. If *C* and *S* are Gaussian random variables with standard deviation *σ* and mean ⟨ *C*⟩ and 0, respectively, then the average cosine of the angle of (*C, S*) is

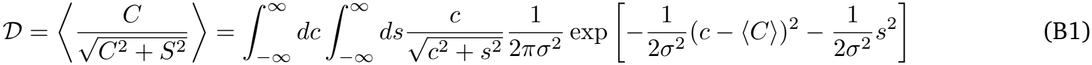

Converting to polar with *c* = *ρ* cos *v* and *s* = *ρ* sin *v*, we get

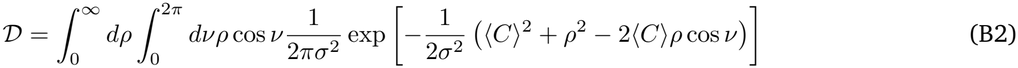

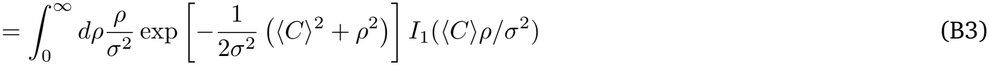

Defining *a* = ⟨*C*⟩*/σ* and *r* = *ρ/σ*, we then get

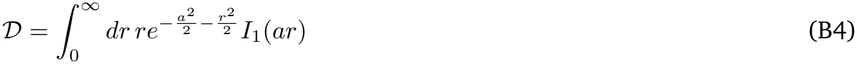

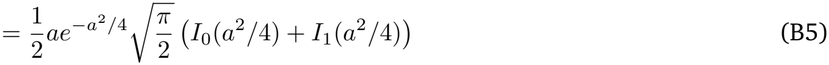

which, using *b* = ⟨*C*⟩^2^*/*4*σ*^2^ = *a*^2^*/*4, gives

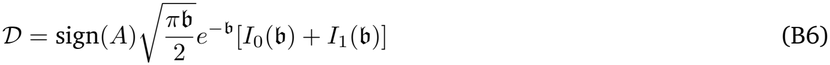

where we have inserted the sign(*A*) term to reintroduce a sign we dropped when we assumed the positive root 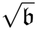. We note that when *µ <* 0, the best estimator of the field orientation is then *ρ* = −(*C, S*), and the directionality would be defined as

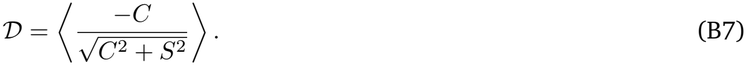

Carrying through the calculation would produce Eq. B5 with the opposite sign. It then makes sense to define the directionality as

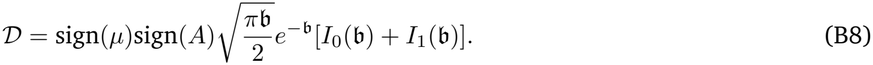

Because of linearity, flipping the sign of *µ* will flip the sign of all values *κ*(*t*) and thus flip the sign of all values of *A*(*t*). Thus, within our paper, these signs will always cancel, and the directionality will be given by Eq. 8. However, if the sign of *A*(*t*) changes without changing *µ* due to, e.g. an AC field or other sign changes of the field, this sign for the directionality will need to be included.

## Appendix C: Generalization to other geometries

### 1. Time dynamics and linear response

We expect our linear response results to hold even for very different geometries than the simple ones we apply here as long as the linear response limit holds. Suppose that we have an arbitrary surface and we write an equation for diffusion and transport on the surface, with **X** being a point on the surface

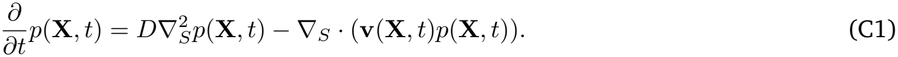

where ^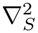^ and ∇_*S*_·· · · indicate the Laplacian and divergence on the surface. If the flow **v**(**X**, *t*) is driven by the electric field, it will be proportional to *µE*_0_, where *E*_0_ is the relevant scale of the applied electric field; we write 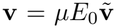. We also rescale our lengths to 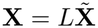, where *L* is some characteristic length - akin to the radius in our circular and spherical models. Given these substitutions,

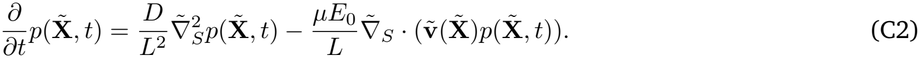

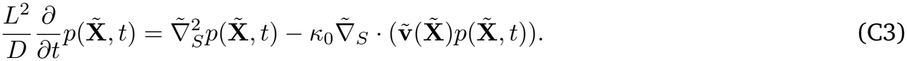

Now, we expand 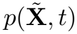 in terms of the eigenfunctions of the surface Laplacian 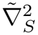. Let’s define 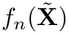 such that 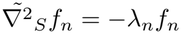, and that *f*_*n*_ are orthonormal, i.e. 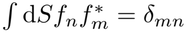. We can then write

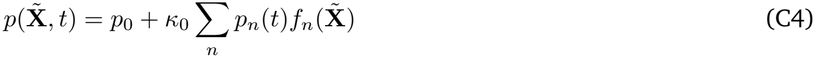

where *p*_*n*_(*t*) are the expansion coefficients, and we’ve separately included the homogeneous solution *p*_0_ which is a constant – so the sum *n* is only over the nonzero eigenvalues.

Plugging this expansion into Eq. C3, we find

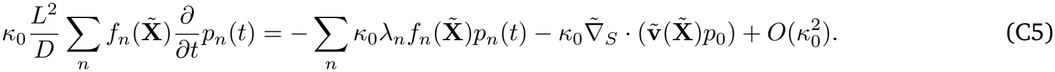

Assuming we are in the linear response regime, we can neglect the final term. If we multiply this equation by 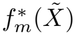 and integrate over the surface, the orthonormality of the eigenfunctions gives us

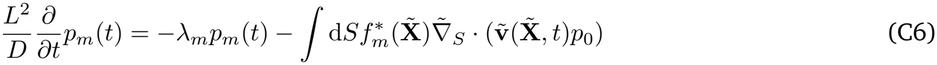

We thus see that this is driving *p*_*m*_ to a target value

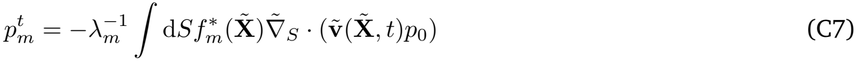

and the relaxation to that value takes the form

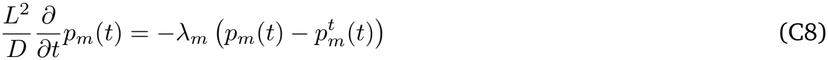

This shows that the relaxation to for mode *m* takes place over a timescale 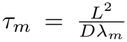. We thus expect that the dynamics of a cell with a more general shape should be limited by the longest timescale – i.e. we expect that we should have

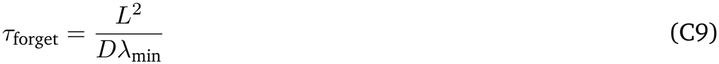

where *λ*_min_ is the smallest nonzero eigenvalue of the surface Laplacian (scaled by *L*^2^).

We can immediately recover results for the circular and spherical cases from this logic. For a sphere with radius *R*, if we choose the length scale *L* = *R*, we are looking for the solution to 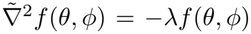. We see that the solution to this equation is the spherical harmonics 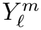, and 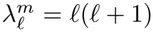 with *l* = 0, 1, 2, · · · [89]. In this case, the lowest-eigenvalue eigenfunction except the trivial constant *l* = 0 is given by *l* = 1, which corresponds to *λ*_min_ = 2. For a circle, the scaled surface Laplacian is just 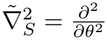, so the eigenfunctions are *e*^*imθ*^ with *m* integer, so the lowest nonconstant eigenfunction corresponds to *m* = 1, so *λ*_min_ = 1.

Our result in Eq. C8 shows that the dynamics of a cell’s response is limited by the longest timescale (smallest nonzero *λ*). However, the timescale of anisotropy arising on a general cell shape could potentially be faster. This could happen if the longest timescale is associated with an eigenmode that does not create a polarization in the direction of the field. For instance, if a cell has its longest extent in the *z* direction, orthogonal to a field in the *x* direction, we would expect the field’s dynamics to be faster than the mode associated with the *z*-direction polarization.

Calculation of these eigenvalues, which are relevant to a huge variety of different problems, including PDE solution and data analysis of surfaces, etc., is possible using finite element methods [40, 90] as well as other approaches [91, 92]. Some analytical results for near-spherical shapes are also known [88].

What are plausible values of the relaxation time for an asymmetric cell? In our experiments, we estimate the HL-60 cells are ~20 microns long, but can have potentially a narrower width and have a rough height of ~5 microns. We calculate eigenvalues and eigenfunctions of the surface Laplacian for a cell modeled as an ellipsoid of length 2, width 2*a*, and height 2*b* using a finite element method [40], specifically using deal.ii [93, 94] and extending the example of [95]. This corresponds to having the length scale *L* = 10 microns; if we choose *a* = 1 and *b* = 1, the shape reduces to a sphere of radius 1 and *λ*_min_ = 2. We thus expect that the relaxation time for mode *m* is 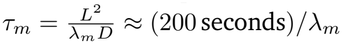 with *L* = 10 microns and *D* = 0.5 microns^2^/s. We see, depending on the range of plausible cell shapes, that the first nonzero eigenvalue is in the rough range of 2.58-3.46, corresponding to a range of ~ 58–78 seconds for the relaxation timescale (Table I). We see that the eigenfunction corresponding to this first mode is generally polarized along the longest axis of the cell, as we expect. As we mentioned above, if the cell’s longest axis is orthogonal to the field, we think the target value for this mode would be small – essentially, the electromigration of sensors would not make this mode large, and the target value for that mode 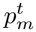 would be small. In this case, the relaxation time would be better approximated by 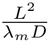 with *m* the mode corresponding to polarization along the direction of the applied field. This is usually the second nonzero eigenvalue, but for very elongated cells (case 1 and 2 in Table I), can be the third nonzero eigenvalue. With this, we expect the relevant eigenvalues to be over the range ~ 6.1 – 11.7 corresponding to a timescale of ~ 17 – 33 seconds (and much shorter for the highly elongated cells where the third eigenvalue should be used). Because real cells prior to the field being turned on may have any angle relative to the field, we would expect the time scale to respond to effectively average over the two relevant time scales of ~ 70 seconds and ~ 20 seconds; this is clearly consistent with the experimental result of *τ*_forget_≈ 35 seconds. However, we emphasize that our range of cell shapes here are fairly artificial compared to the complex cell shapes of HL60 [65] and the estimates of cell shape in the experiment are rough. Another potential issue that we raised in the main text is that the measurement of the diffusion coefficient *D* was done on immobilized, latrunculin-treated cells – so the diffusion coefficient in crawling cells may vary slightly.

**TABLE I.**
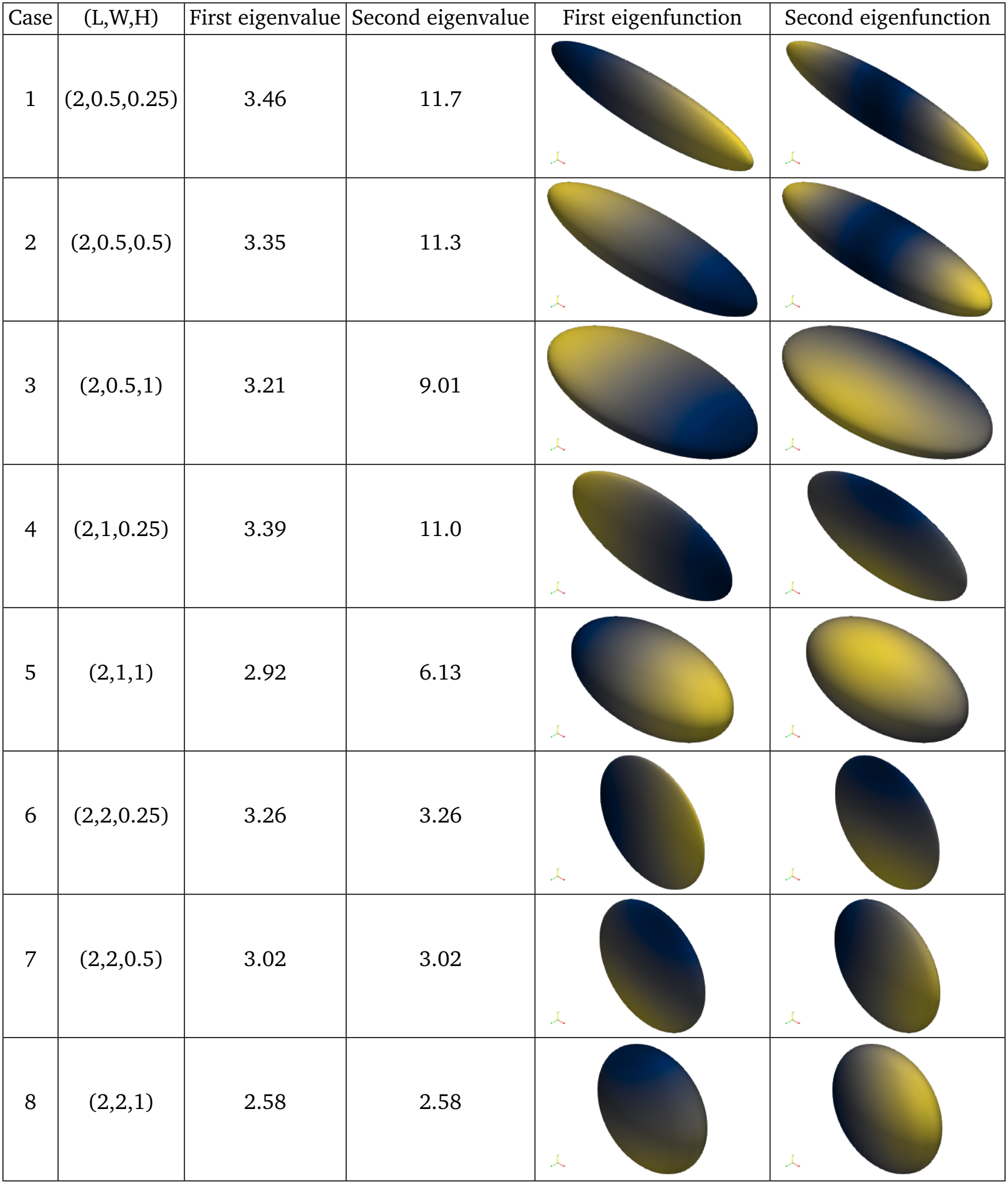
First and second nonzero eigenvalues for cells of a range of shapes, along with corresponding eigenfunctions. Units are chosen so that the longest axis has length 2, so the first nonzero eigenvalue for a sphere is 2 (see text). Color indicates the value of the eigenfunction, with yellow indicating positive values, though the overall sign is arbitrary. First eigenfunctions normally switch sign along the longest axis, indicating polarization of the sensor along the long axis – akin to our cos *θ* in 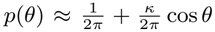. Second eigenfunctions are often indicating polarization along an orthogonal axis to the long axis, though for cases 1 and 2, they indicate a higher-mode polarization along the long axis.

| Case | (L,W,H) | First eigenvalue | Second eigenvalue | First eigenfunction | Second eigenfunction |
|---|---|---|---|---|---|
| 1 | (2,0.5,0.25) | 3.46 | 11.7 |  |  |
| 2 | (2,0.5,0.5) | 3.35 | 11.3 |  |  |
| 3 | (2,0.5,1) | 3.21 | 9.01 |  |  |
| 4 | (2,1,0.25) | 3.39 | 11.0 |  |  |
| 5 | (2,1,1) | 2.92 | 6.13 |  |  |
| 6 | (2,2,0.25) | 3.26 | 3.26 |  |  |
| 7 | (2,2,0.5) | 3.02 | 3.02 |  |  |
| 8 | (2,2,1) | 2.58 | 2.58 |  |  |

### 2. Directionality for a spherical cell

While we have not been able to provide results for how directionality depends on anisotropy for an arbitrary surface (see, e.g. the complications arising in [33]), we can generalize the result of Eq. 8 to the case of a spherical cell in three dimensions.

For a spherical cell, sensors in a constant *E* field are distributed with a von Mises-Fisher distribution [32], *p*(*θ, ϕ*) ~*e*^*κÊ·û*^, where *Ê* is a unit vector in the direction of the E field and *û* = (cos *ϕ* sin *θ*, sin *ϕ* sin *θ*, cos *θ*). Assuming the E field is in the *x* direction, this is simply *p*(*θ, ϕ*) ~ *e*^*κ* cos *ϕ* sin *θ*^. Then to linear order in *κ*,

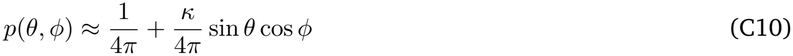

In 2D, we defined anisotropy as 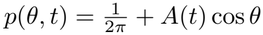 in 3D we choose

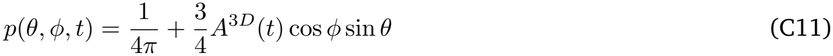

This somewhat odd factor of 3/4 is a convention to ensure that the expectation ⟨cos *ϕ* sin *θ*⟩ = *πA*^3*D*^ in 3D, just as ⟨cos *θ*⟩ = *Aπ* in 2D, which we will see below will simplify our analysis. If the cell has *N* sensors at angular positions (*θ*_*i*_, *ϕ*_*i*_), the maximum-likelihood estimator for the field direction, assuming that the cell is on a substrate in the *xy* plane and that the field is parallel to this substrate, is that the field orientation is in the direction of

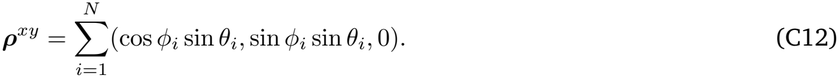

As in Appendix A, we can then work out the mean and standard deviation of *C* = ∑ _*i*_ cos *ϕ*_*i*_ sin *θ*_*i*_ and *S* = ∑ _*i*_ sin *ϕ*_*i*_ sin *θ*_*i*_ under the assumption that we have the linear form of Eq. C11, finding

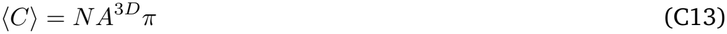

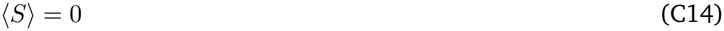

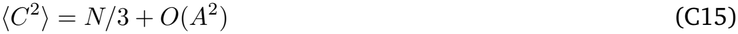

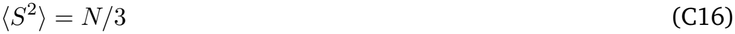

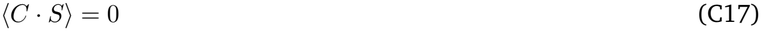

We can then see that *C* and *S* are sums of independent identically distributed variables – so we expect them to be Gaussian with variance *σ*^2^ = *N/*3 (up to linear order in *A*). This means that we expect in 3D that directionality will be given by Eq. 8 but with 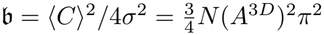.

We also note that if we use the linear form of Eq. C11, we should adjust the formula for the cathode-to-anode ratio of fluorescence. This would become

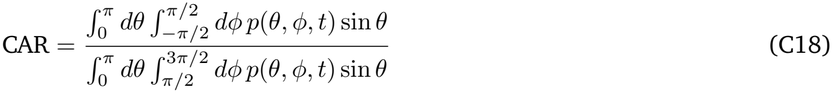

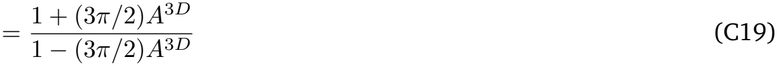

i.e., the result for the 2D system but with a factor of 3*π/*2 ≈ 4.7 instead of 4.

## Appendix D: Effect of *D*_max_, geometry, and CAR assumptions on fitting

### 1. Effect of *D*_max_

We show in this section some analysis of how choosing *D*_max_ in the analysis of Fig. 4 changes our results. Because of the form of Eq. 12, it would be possible to decrease directionality either by decreasing *N* or decreasing *D*_max_. As a result, a choice of this parameter can systematically, though generally weakly, affect the estimated value of the effective number of sensors *N*_eff_. In the main text we chose *D*_max_ = 1. In Fig. S1, we vary *D*_max_ and repeat the analysis of Fig. 4. The lowest possible value of *D*_max_ is ≈ 0.4, as we see the directionality reaches this value in Fig. 4. As *D*_max_ decreases toward the lowest possible value, the estimated number of sensors increases, with increasingly broad distributions of fit values as we are constraining the fit to implausibly small values of *D*_max_. As long as *D*_max_ is not fine-tuned to 0.4, the effective number of sensors is likely no more than 100 or so. For a spherical cell with radius ~ 10 microns, this would be ~ 0.08 sensors per square micron – still likely low enough density for sensors to be resolved as diffraction-limited spots. We also suspect that the maximum directionality for neutrophils in the under-agarose experiments shown in Fig. 4 is significantly higher than 0.4. While it is difficult to generalize between 2D and 3D migration [96], earlier measurements by Belliveau et al. showed that increasing electric fields increased the directionality measured in 3D collagen matrices increases up to at least ~ 0.8 (personal communication), suggesting *D*_max_ is not small enough to change our results significantly.

**FIG. S1.**
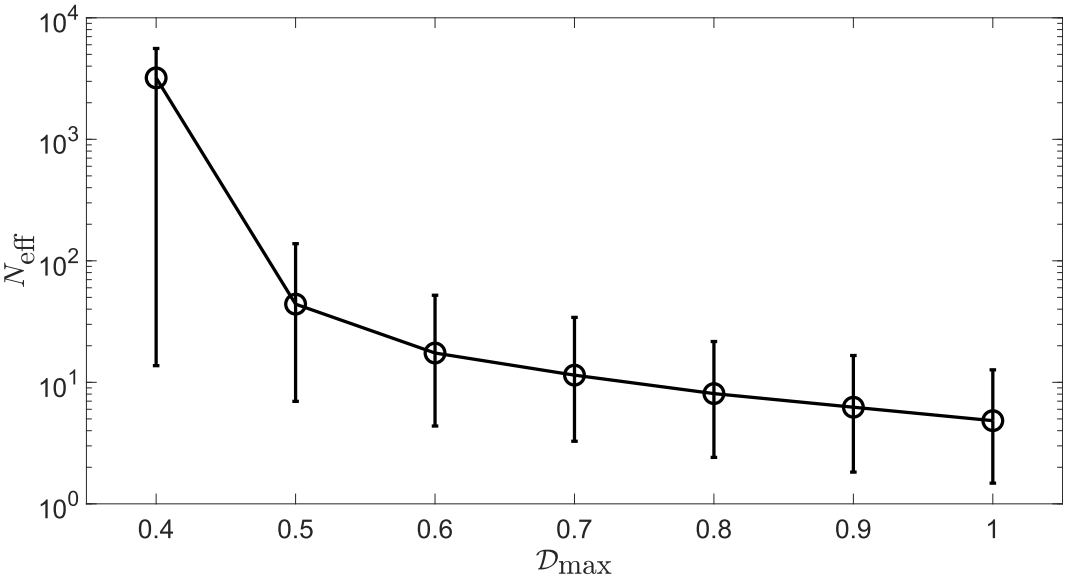
The best fit values of *N*_eff_ vary systematically with the assumed value of *D*_max_. Error bars contain 95% of bootstraps.

### 2. Alternate fitting assumptions for CAR and directionality

If we fit the results of Fig. 4 assuming a three-dimensional cell shape, as in Section C 2, we find very similar fit parameters to those of Fig. 4, *τ*_forget_ = 37 ±10 s, *κ* =− 0.59 ±0.11, and *N*_eff_ = 4.5 ±2.8. Different assumptions for the cell shape or slight alternate definitions of the cathode-to-anode ratio would likely also change the fit values of *κ* and *N* slightly. We see for a sphere

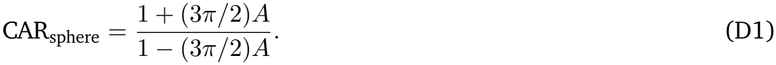

Within the main paper, we have used (1 + 4*A*)*/*(1 4*A*) for a circle, but of course the HL60s are not perfectly circular or spherical. Another possible issue is that, due to some experimental imaging and segmentation reasons, the cathodal fluorescence and anodal fluorescence are quantified as in [10] – over the range of 120 to 240 degrees for the anode and 60 to 60 degrees for the cathode. Given this, if you compute the CAR as 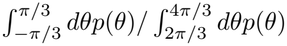, we get

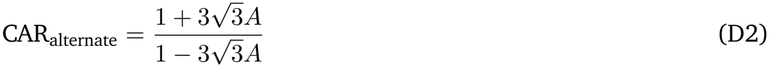

Using this alternate form, we find *τ*_forget_ = 37.5 ± 9.7 s, *κ* = −0.36± 0.07, and *N*_eff_ = 8.1 ± 5. Other subtleties, such as modeling the effects of different imaging approaches etc., could also change the prefactor in front of *A* in our formulas for CAR, and would correspondingly slightly shift the values of *κ* and *N*_eff_ as well.

## Appendix E: Oscillating field orientation

It’s straightforward to expand our linear response results to a case where the field orientation *ψ* is time-dependent as well. In this case, the Smoluchowski equation for the sensor positions is

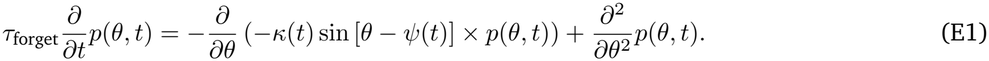

For oscillating field orientation, we expect the linear response will have pieces proportional to cos *θ* and sin *θ* (or equivalently a time-varying phase), so we use the Ansatz

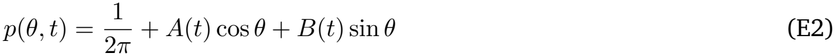

where *A* and *B* are both linear in the maximum *κ, κ*_0_, which gives us

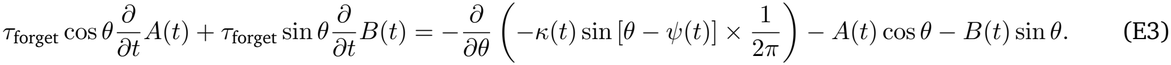

where we have neglected the nonlinear terms proportional to *Aκ* and *Bκ*. We then get, evaluating the derivative and applying angle addition formulas

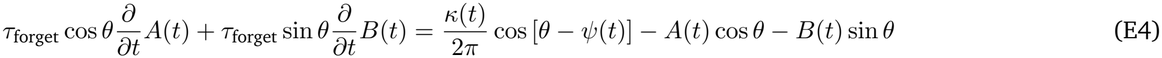

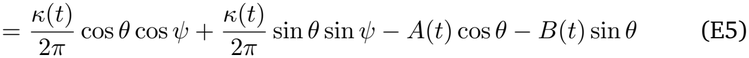

Equating the coefficients of cos *θ* and sin *θ* we find

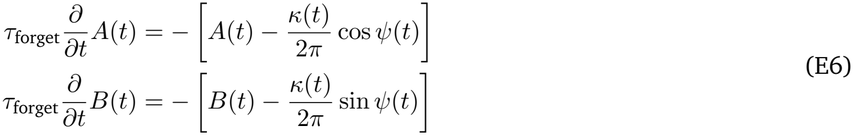

This has some simple consequences. For instance, suppose that the field direction is switched between +*δ* and ™*δ* periodically, but the magnitude kept constant, as was done, e.g. by. Experimentally, cells are found to move in the vector average of the two applied field directions if the field is switched rapidly – so we would expect that the cell would on average have an anisotropy solely in the *x* direction, i.e. 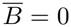 but *Ā* ≠ 0. If *A*(*t*) and *B*(*t*) have period *T*, we can evaluate the integral of Eq. E6 over one period. In this case, 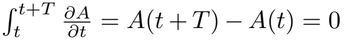, and similarly for *B*. This shows that the time average 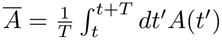 obeys

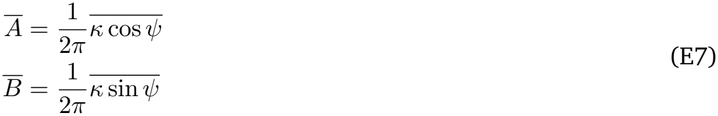

If the field is being switched between *δ* and *δ* with constant magnitude (i.e. constant *κ*), we see 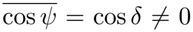 but 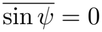. This shows, as we expect, that on average, the cell is only polarized along the *x* axis.

We note this averaging result is not unique to our linear response model – we also found something very similar with a different phenomenological model [12] – we expect that something like this has to be true of many models that have an underlying cell polarity reoriented by the field.

## Appendix F: Jensen’s inequality and time-averaged directionality

Jensen’s inequality states that for a *convex* function *ϕ* that

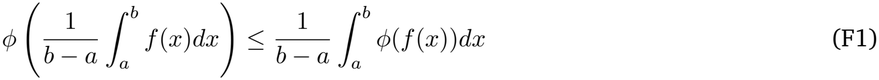

or – because if *ϕ* is convex, then Φ = −*ϕ* is concave and vice versa– for a concave function Φ,

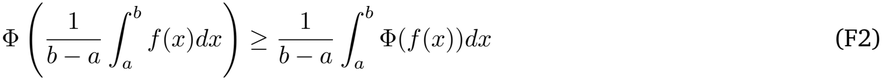

For our functions with period *T*, it makes sense to define a time-average as

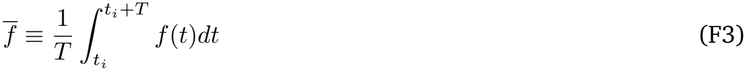

with *t*_*i*_ some arbitrary initial time.

Jensen’s inequality then states for a concave function Φ that

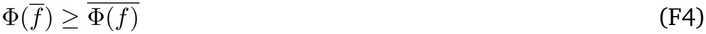

The directionality *D* is a function of the anisotropy *A*(*t*),

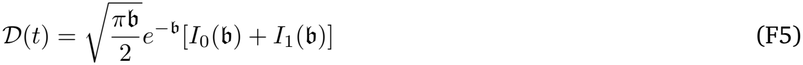

with *b*(*t*) = *NA*(*t*)^2^*π*^2^*/*2. This function (*A*) is concave in *A* as long as *A* does not change sign, e.g. *A >* 0 for *µ >* 0 or *A <* 0 for *µ <* 0; (see Appendix B) – it has negative second derivative. Jensen’s inequality thus shows that

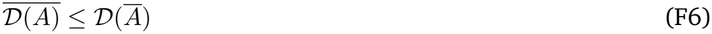

We showed in the main text (Eq. 18) that 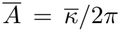, so evaluating the directionality at the time-average of the anisotropy is just the same as taking the time-average of the applied electric field. This shows

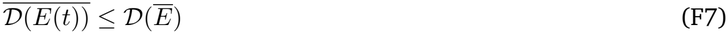

as we claimed in the main text.

**FIG. S2.**
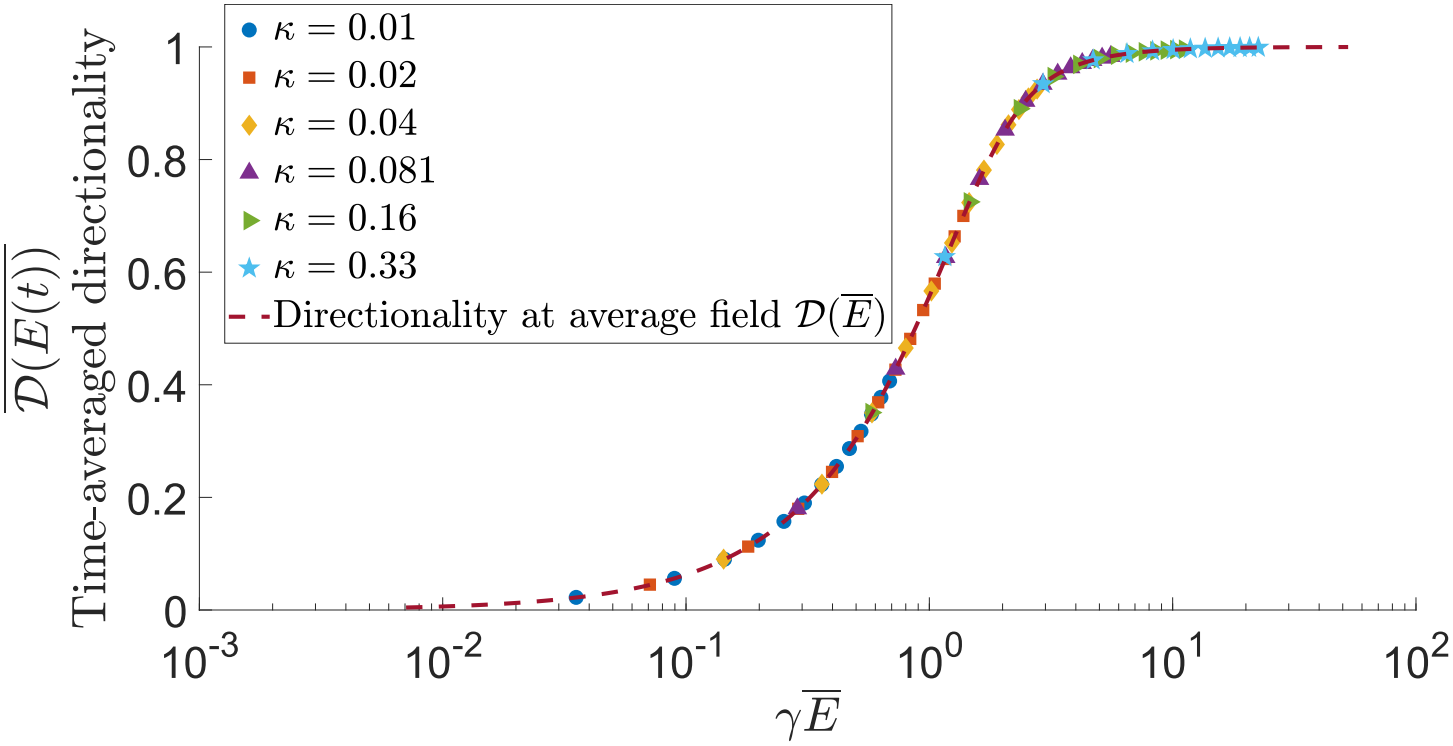
Collapse of time-averaged directionality 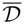 as a function of *γ Ē* for pulsed fields. This figure is generated in the same way and with the same parameters as Fig. 7, with the exception of period *T* = 0.1*τ*_forget_, maximum time 10*τ*_forget_, and timestep Δ*t* = 10^*−*4^*τ*_forget_.

## Notes

### Competing Interest Statement

The authors have declared no competing interest.

### Summary of Updates

Clarifications and new discussion of shot noise limitation and limitations of assumptions of fits; added new figure panel on viscosity effects; other minor fixes and clarifications

## References

[1] Colin D McCaig, Ann M Rajnicek, Bing Song, and Min Zhao. Controlling cell behavior electrically: current views and future potential. Physiological reviews, 2005.

[2] Min Zhao. Electrical fields in wound healing—an overriding signal that directs cell migration. In Seminars in Cell & Developmental Biology, volume 20, pages 674–682. Elsevier, 2009.

[3] Shuvasree SenGupta, Carole A Parent, and James E Bear. The principles of directed cell migration. Nature Reviews Molecular Cell Biology, pages 1–19, 2021.

[4] Greg M Allen, Alex Mogilner, and Julie A Theriot. Electrophoresis of cellular membrane components creates the directional cue guiding keratocyte galvanotaxis. Current Biology, 23(7):560–568, 2013.

[5] Anyesha Sarkar, Brian M Kobylkevich, David M Graham, and Mark A Messerli. Electromigration of cell surface macromolecules in DC electric fields during cell polarization and galvanotaxis. Journal of theoretical biology, 478:58–73, 2019.

[6] Brian M Kobylkevich, Anyesha Sarkar, Brady R Carlberg, Ling Huang, Suman Ranjit, David M Graham, and Mark A Messerli. Reversing the direction of galvanotaxis with controlled increases in boundary layer viscosity. Physical Biology, 15(3):036005, 2018.

[7] Martin J Brown and Leslie M Loew. Electric field-directed fibroblast locomotion involves cell surface molecular reorganization and is calcium independent. The Journal of Cell Biology, 127(1):117–128, 1994.

[8] S McLaughlin and MM Poo. The role of electro-osmosis in the electric-field-induced movement of charged macromolecules on the surfaces of cells. Biophysical Journal, 34(1):85–93, 1981.

[9] Slawomir Lasota, Eliza Zimolag, Sylwia Bobis-Wozowicz, Jagoda Pilipiuk, and Zbigniew Madeja. The dynamics of the electrotactic reaction of mouse 3T3 fibroblasts. Biochimica et Biophysica Acta (BBA)-Molecular Cell Research, 1871(2):119647, 2024.

[10] Nathan M Belliveau, Matthew J Footer, Amy Platenkamp, Celeste Rodriguez, Heonsu Kim, Christopher K Prinz, Aaron P van Loon, Yubin Lin, Tara E Eustis, Michelle M Chan, et al. Galvanin (tmem154) is an electric-field sensor for directed cell migration. Cell, 189(13):4107–4121, 2026.

[11] Hans Gruler and Richard Nuccitelli. The galvanotaxis response mechanism of keratinocytes can be modeled as a proportional controller. Cell biochemistry and biophysics, 33:33–51, 2000.

[12] Ifunanya Nwogbaga and Brian A Camley. Coupling cell shape and velocity leads to oscillation and circling in keratocyte galvanotaxis. Biophysical Journal, 122(1):130–142, 2023.

[13] Thomas P Prescott, Kan Zhu, Min Zhao, and Ruth E Baker. Quantifying the impact of electric fields on single-cell motility. Biophysical Journal, 120:3363–3373, 2021.

[14] Danny Fuller, Wen Chen, Micha Adler, Alex Groisman, Herbert Levine, Wouter-Jan Rappel, and William F Loomis. External and internal constraints on eukaryotic chemotaxis. Proceedings of the National Academy of Sciences, 107(21):9656–9659, 2010.

[15] Igor Segota, Surin Mong, Eitan Neidich, Archana Rachakonda, Catherine J Lussenhop, and Carl Franck. High fidelity information processing in folic acid chemotaxis of Dictyostelium amoebae. Journal of The Royal Society Interface, 10(88):20130606, 2013.

[16] Kazunari Kaizu, Wiet De Ronde, Joris Paijmans, Koichi Takahashi, Filipe Tostevin, and Pieter Rein Ten Wolde. The Berg-Purcell limit revisited. Biophysical Journal, 106(4):976–985, 2014.

[17] Howard C Berg and Edward M Purcell. Physics of chemoreception. Biophysical Journal, 20(2):193–219, 1977.

[18] Pieter Rein ten Wolde, Nils B Becker, Thomas E Ouldridge, and Andrew Mugler. Fundamental limits to cellular sensing. Journal of Statistical Physics, 162:1395–1424, 2016.

[19] Herbert Levine and Wouter-Jan Rappel. The physics of eukaryotic chemotaxis. Physics today, 66(2):24–30, 2013.

[20] William Bialek and Sima Setayeshgar. Physical limits to biochemical signaling. Proceedings of the National Academy of Sciences, 102(29):10040–10045, 2005.

[21] Henry H Mattingly, Keita Kamino, Jude Ong, Rafaela Kottou, Thierry Emonet, and Benjamin B Machta. E. coli chemosensing accuracy is not limited by stochastic molecule arrivals. Nature Physics, pages 1–8, 2026.

[22] Aparajita Kashyap, Wei Wang, and Brian A Camley. Trade-offs in concentration sensing in dynamic environments. Biophysical Journal, 123(10):1184–1194, 2024.

[23] Andrew J Bernoff, Alexandra Jilkine, Adrián Navarro Hernández, and Alan E Lindsay. Single-cell directional sensing from just a few receptor binding events. Biophysical Journal, 122(15):3108–3116, 2023.

[24] Sean D Lawley, Alan E Lindsay, and Christopher E Miles. Receptor organization determines the limits of single-cell source location detection. Physical Review Letters, 125(1):018102, 2020.

[25] Daiqiu Mou and Yuansheng Cao. Optimal cell shape for accurate chemical gradient sensing in eukaryote chemotaxis. Physical Review Research, 7(3):033001, 2025.

[26] William Bialek. Biophysics: searching for principles. Princeton University Press, 2012.

[27] Wei Wang and Brian A Camley. Limits on the accuracy of contact inhibition of locomotion. Physical Review E, 109(5):054408, 2024.

[28] Motasem ElGamel and Andrew Mugler. Effects of molecular noise on cell size control. Physical Review Letters, 132(9):098403, 2024.

[29] Indrajit Badvaram and Brian A Camley. Physical limits to membrane curvature sensing by a single protein. Physical Review E, 108(6):064407, 2023.

[30] Eric D Siggia and Massimo Vergassola. Decisions on the fly in cellular sensory systems. Proceedings of the National Academy of Sciences, 110(39):E3704–E3712, 2013.

[31] Jonathan Desponds, Massimo Vergassola, and Aleksandra M Walczak. A mechanism for hunchback promoters to readout morphogenetic positional information in less than a minute. Elife, 9:e49758, 2020.

[32] Ifunanya Nwogbaga, A Hyun Kim, and Brian A Camley. Physical limits on galvanotaxis. Physical Review E, 108(6):064411, 2023.

[33] Ifunanya Nwogbaga and Brian A Camley. Cell shape and orientation control galvanotactic accuracy. Soft Matter, 20(44):8866–8887, 2024.

[34] David Chandler. Introduction to Modern Statistical Mechanics. Oxford University Press, Oxford, UK, 1987.

[35] Mu-ming Poo. In situ electrophoresis of membrane components. Annual review of biophysics and bioengineering, 10(1):245–276, 1981.

[36] Robert G Endres and Ned S Wingreen. Accuracy of direct gradient sensing by single cells. Proceedings of the National Academy of Sciences, 105(41):15749–15754, 2008.

[37] Masahiro Ueda and Tatsuo Shibata. Stochastic signal processing and transduction in chemotactic response of eukaryotic cells. Biophysical Journal, 93(1):11–20, 2007.

[38] Greg M Allen, Kun Chun Lee, Erin L Barnhart, Mark A Tsuchida, Cyrus A Wilson, Edgar Gutierrez, Alexander Groisman, Julie A Theriot, and Alex Mogilner. Cell mechanics at the rear act to steer the direction of cell migration. Cell Systems, 11(3):286–299, 2020.

[39] K Franke and H Gruler. Galvanotaxis of human granulocytes: electric field jump studies. European Biophysics Journal, 18:334–346, 1990.

[40] Martin Reuter, Silvia Biasotti, Daniela Giorgi, Giuseppe Patanè, and Michela Spagnuolo. Discrete Laplace–Beltrami operators for shape analysis and segmentation. Computers & Graphics, 33(3):381–390, 2009.

[41] Diego Krapf. Mechanisms underlying anomalous diffusion in the plasma membrane. Current topics in membranes, 75:167–207, 2015.

[42] Kripa Gowrishankar, Subhasri Ghosh, Suvrajit Saha, C Rumamol, Satyajit Mayor, and Madan Rao. Active remodeling of cortical actin regulates spatiotemporal organization of cell surface molecules. Cell, 149(6):1353–1367, 2012.

[43] Nathan W Goehring, Philipp Khuc Trong, Justin S Bois, Debanjan Chowdhury, Ernesto M Nicola, Anthony A Hyman, and Stephan W Grill. Polarization of PAR proteins by advective triggering of a pattern-forming system. Science, 334(6059):1137–1141, 2011.

[44] Brian A Camley, Yunsong Zhang, Yanxiang Zhao, Bo Li, Eshel Ben-Jacob, Herbert Levine, and Wouter-Jan Rappel. Polarity mechanisms such as contact inhibition of locomotion regulate persistent rotational motion of mammalian cells on micropatterns. Proceedings of the National Academy of Sciences, 111(41):14770–14775, 2014.

[45] Yuan Xiong, Chuan-Hsiang Huang, Pablo A Iglesias, and Peter N Devreotes. Cells navigate with a local-excitation, global-inhibition-biased excitable network. Proceedings of the National Academy of Sciences, 107(40):17079–17086, 2010.

[46] Peter N Devreotes, Sayak Bhattacharya, Marc Edwards, Pablo A Iglesias, Thomas Lampert, and Yuchuan Miao. Excitable signal transduction networks in directed cell migration. Annual Review of Cell and Developmental Biology, 33(1):103–125, 2017.

[47] Shardool Kulkarni, Francesc Tebar, Carles Rentero, Min Zhao, and Pablo Saez. Competing signaling pathways controls electrotaxis. iScience, 28(5), 2025.

[48] Eugene P Petrov, Rafayel Petrosyan, and Petra Schwille. Translational and rotational diffusion of micrometer-sized solid domains in lipid membranes. Soft Matter, 8(29):7552–7555, 2012.

[49] Eugene P Petrov and Petra Schwille. Translational diffusion in lipid membranes beyond the Saffman-Delbrück approximation. Biophysical Journal, 94(5):L41–L43, 2008.

[50] PG Saffman and M Delbrück. Brownian motion in biological membranes. Proceedings of the National Academy of Sciences, 72(8):3111–3113, 1975.

[51] BD Hughes, BA Pailthorpe, and LR White. The translational and rotational drag on a cylinder moving in a membrane. Journal of Fluid Mechanics, 110:349–372, 1981.

[52] Brian A Camley, Michael G Lerner, Richard W Pastor, and Frank LH Brown. Strong influence of periodic boundary conditions on lateral diffusion in lipid bilayer membranes. The Journal of chemical physics, 143(24), 2015.

[53] Naomi Oppenheimer and Haim Diamant. Correlated diffusion of membrane proteins and their effect on membrane viscosity. Biophysical Journal, 96(8):3041–3049, 2009.

[54] Brian A Camley and Frank LH Brown. Motion of objects embedded in lipid bilayer membranes: Advection and effective viscosity. The Journal of Chemical Physics, 151(12), 2019.

[55] Naomi Oppenheimer and Haim Diamant. In-plane dynamics of membranes with immobile inclusions. Physical review letters, 107(25):258102, 2011.

[56] Mark L Henle and Alex J Levine. Hydrodynamics in curved membranes: The effect of geometry on particulate mobility. Physical Review E—Statistical, Nonlinear, and Soft Matter Physics, 81(1):011905, 2010.

[57] Brian A Camley and Frank LH Brown. Beyond the creeping viscous flow limit for lipid bilayer membranes: Theory of single-particle microrheology, domain flicker spectroscopy, and long-time tails. Physical Review E—Statistical, Nonlinear, and Soft Matter Physics, 84(2):021904, 2011.

[58] Kinneret Keren, Zachary Pincus, Greg M Allen, Erin L Barnhart, Gerard Marriott, Alex Mogilner, and Julie A Theriot. Mechanism of shape determination in motile cells. Nature, 453(7194):475–480, 2008.

[59] Matthew Pittman, Ernest Iu, Keva Li, Junjie Chen, Sergey Potnikov, and Yun Chen. Membrane ruffling is a mechanosensor of extracellular fluid viscosity. Biophysical Journal, 121(3):267a, 2022.

[60] Kaustav Bera, Alexander Kiepas, Inês Godet, Yizeng Li, Pranav Mehta, Brent Ifemembi, Colin D Paul, Anindya Sen, Selma A Serra, Konstantin Stoletov, et al. Extracellular fluid viscosity enhances cell migration and cancer dissemination. Nature, 611(7935):365–373, 2022.

[61] Chiranjeevi Korupalli, Hui Li, Nhien Nguyen, Fwu-Long Mi, Yen Chang, Yu-Jung Lin, and Hsing-Wen Sung. Conductive materials for healing wounds: their incorporation in electroactive wound dressings, characterization, and perspectives. Advanced healthcare materials, 10(6):2001384, 2021.

[62] Joseph C Ojingwa and R Rivkah Isseroff. Electrical stimulation of wound healing. Journal of Investigative Dermatology, pages 1–12, 2003.

[63] Xi Ren, Huanbo Sun, Jie Liu, Xiaowei Guo, Jingzhuo Huang, Xupin Jiang, Yiming Zhang, Yuesheng Huang, Dongli Fan, and Jiaping Zhang. Keratinocyte electrotaxis induced by physiological pulsed direct current electric fields. Bioelectrochemistry, 127:113–124, 2019.

[64] Samuel J Lord, Katrina B Velle, R Dyche Mullins, and Lillian K Fritz-Laylin. Superplots: Communicating reproducibility and variability in cell biology. Journal of Cell Biology, 219(6), 2020.

[65] Lillian K Fritz-Laylin, Megan Riel-Mehan, Bi-Chang Chen, Samuel J Lord, Thomas D Goddard, Thomas E Ferrin, Susan M Nicholson-Dykstra, Henry Higgs, Graham T Johnson, Eric Betzig, et al. Actin-based protrusions of migrating neutrophils are intrinsically lamellar and facilitate direction changes. eLife, 6:e26990, 2017.

[66] Georg R Walther, Athanasius FM Marée, Leah Edelstein-Keshet, and Verônica A Grieneisen. Deterministic versus stochastic cell polarisation through wave-pinning. Bulletin of Mathematical Biology, 74(11):2570–2599, 2012.

[67] Mariia Kryvoruchko and Brian A Camley. Intrinsic stochasticity in cell polarity and contact inhibition of locomotion. arXiv preprint arXiv:2604.18851, 2026.

[68] Inbal Hecht, Monica L Skoge, Pascale G Charest, Eshel Ben-Jacob, Richard A Firtel, William F Loomis, Herbert Levine, and Wouter-Jan Rappel. Activated membrane patches guide chemotactic cell motility. PLoS Computational Biology, 7(6):e1002044, 2011.

[69] Matthew P Neilson, Douwe M Veltman, Peter JM van Haastert, Steven D Webb, John A Mackenzie, and Robert H Insall. Chemotaxis: a feedback-based computational model robustly predicts multiple aspects of real cell behaviour. PLoS biology, 9(5):e1000618, 2011.

[70] Changji Shi, Chuan-Hsiang Huang, Peter N Devreotes, and Pablo A Iglesias. Interaction of motility, directional sensing, and polarity modules recreates the behaviors of chemotaxing cells. PLoS Computational Biology, 9(7):e1003122, 2013.

[71] Adrian Moure and Hector Gomez. Computational model for amoeboid motion: Coupling membrane and cytosol dynamics. Physical Review E, 94(4):042423, 2016.

[72] Bo-jian Lin, Shun-hao Tsao, Alex Chen, Shu-Kai Hu, Ling Chao, and Pen-hsiu Grace Chao. Lipid rafts sense and direct electric field-induced migration. Proceedings of the National Academy of Sciences, 114(32):8568–8573, 2017.

[73] Alexandra Jilkine and Leah Edelstein-Keshet. A comparison of mathematical models for polarization of single eukaryotic cells in response to guided cues. PLoS computational biology, 7(4):e1001121, 2011.

[74] Wouter-Jan Rappel and Leah Edelstein-Keshet. Mechanisms of cell polarization. Current opinion in systems biology, 3:43–53, 2017.

[75] Monica Skoge, Haicen Yue, Michael Erickstad, Albert Bae, Herbert Levine, Alex Groisman, William F Loomis, and Wouter-Jan Rappel. Cellular memory in eukaryotic chemotaxis. Proceedings of the National Academy of Sciences, 111(40):14448–14453, 2014.

[76] Harrison V Prentice-Mott, Yasmine Meroz, Andreas Carlson, Michael A Levine, Michael W Davidson, Daniel Irimia, Guillaume T Charras, Lakshminarayanan Mahadevan, and Jagesh V Shah. Directional memory arises from long-lived cytoskeletal asymmetries in polarized chemotactic cells. Proceedings of the National Academy of Sciences, 113(5):1267–1272, 2016.

[77] Richa Karmakar, Man-Ho Tang, Haicen Yue, Daniel Lombardo, Aravind Karanam, Brian A Camley, Alex Groisman, and Wouter-Jan Rappel. Cellular memory in eukaryotic chemotaxis depends on the background chemoattractant concentration. Physical Review E, 103(1):012402, 2021.

[78] Brian A Camley and Wouter-Jan Rappel. Velocity alignment leads to high persistence in confined cells. Physical Review E, 89(6):062705, 2014.

[79] Isabela Corina Fortunato, David B Brückner, Steffen Grosser, Rohit Nautiyal, Leone Rossetti, Miquel Bosch-Padrós, Jonel Trebicka, Pere Roca-Cusachs, Raimon Sunyer, Edouard Hannezo, et al. Single-cell migration along and against confined haptotactic gradients. Nature Physics, 21(10):1638–1647, 2025.

[80] Bashar Hamza, Elisabeth Wong, Sachin Patel, Hansang Cho, Joseph Martel, and Daniel Irimia. Retrotaxis of human neutrophils during mechanical confinement inside microfluidic channels. Integrative Biology, 6(2):175–183, 2014.

[81] Yaohui Sun, Brian Reid, Yan Zhang, Kan Zhu, Fernando Ferreira, Alejandro Estrada, Yuxin Sun, Bruce W Draper, Haicen Yue, Calina Copos, et al. Electric field–guided collective motility initiation of large epidermal cell groups. Molecular biology of the cell, 34(5):ar48, 2023.

[82] Kurmanbek Kaiyrbekov and Brian A Camley. Does nematic order allow groups of elongated cells to sense electric fields better? Plos one, 20(6):e0325800, 2025.

[83] Gawoon Shim, Danelle Devenport, and Daniel J Cohen. Overriding native cell coordination enhances external programming of collective cell migration. Proceedings of the National Academy of Sciences, 118(29), 2021.

[84] Li Li, Robert Hartley, Bjoern Reiss, Yaohui Sun, Jin Pu, Dan Wu, Francis Lin, Trung Hoang, Soichiro Yamada, Jianxin Jiang, et al. E-cadherin plays an essential role in collective directional migration of large epithelial sheets. Cellular and Molecular Life Sciences, 69(16):2779–2789, 2012.

[85] Fernando Ferreira, Sofia Moreira, Min Zhao, and Elias H Barriga. Stretch-induced endogenous electric fields drive directed collective cell migration in vivo. Nature Materials, pages 1–9, 2025.

[86] Brian Camley, Juliane Zimmermann, Herbert Levine, and Wouter-Jan Rappel. Emergent collective chemotaxis without single-cell gradient sensing. Physical Review Letters, 116:098101, 2016.

[87] Brian A Camley. Collective gradient sensing and chemotaxis: modeling and recent developments. Journal of Physics: Condensed Matter, 30(22):223001, 2018.

[88] Suresh Eswarathasan and Theodore Kolokolnikov. Laplace–Beltrami spectrum of ellipsoids that are close to spheres and analytic perturbation theory. IMA Journal of Applied Mathematics, 87(1):20–49, 2022.

[89] George B Arfken, Hans J Weber, and Frank E Harris. Mathematical methods for physicists: a comprehensive guide. Academic press, 2011.

[90] Martin Reuter, Franz-Erich Wolter, and Niklas Peinecke. Laplace–Beltrami spectra as ‘Shape-DNA’of surfaces and solids. Computer-Aided Design, 38(4):342–366, 2006.

[91] Daniel R Venn and Steven J Ruuth. A meshfree method for eigenvalues of differential operators on surfaces, including Steklov problems. arXiv preprint arXiv:2410.04336, 2024.

[92] Yingying Wu, Tianqi Wu, and Shing-Tung Yau. Surface eigenvalues with lattice-based approximation in comparison with analytical solution. arXiv preprint arXiv:2203.03603, 2022.

[93] Pasquale C. Africa, Daniel Arndt, Wolfgang Bangerth, Bruno Blais, Marc Fehling, Rene Gassmöller, Timo Heister, Luca Heltai, Sebastian Kinnewig, Martin Kronbichler, Matthias Maier, Peter Munch, Magdalena Schreter-Fleischhacker, Jan P. Thiele, Bruno Turcksin, David Wells, and Vladimir Yushutin. The deal.II library, version 9.6. Journal of Numerical Mathematics, 32(4):369–380, 2024.

[94] Daniel Arndt, Wolfgang Bangerth, Denis Davydov, Timo Heister, Luca Heltai, Martin Kronbichler, Matthias Maier, Jean-Paul Pelteret, Bruno Turcksin, and David Wells. The deal.II finite element library: Design, features, and insights. Computers & Mathematics with Applications, 81:407–422, 2021.

[95] Toby D. Young and Wolfgang Bangerth. The step-36 tutorial program, step 36. https://www.dealii.org/current/doxygen/deal.II/step_36.html. Accessed: 2021-29-01.

[96] Guillaume Charras and Erik Sahai. Physical influences of the extracellular environment on cell migration. Nature reviews Molecular cell biology, 15(12):813–824, 2014.

[97] Tom J Zajdel, Gawoon Shim, Linus Wang, Alejandro Rossello-Martinez, and Daniel J Cohen. SCHEEPDOG: programming electric cues to dynamically herd large-scale cell migration. Cell Systems, 10(6):506–514, 2020.

